# Therapy-driven clonal selection links genomic heterogeneity to transcriptional cell-state adaptation in small cell lung cancer

**DOI:** 10.64898/2026.11.26.769450

**Authors:** Laura C Kaiser, Wassim Jaoui, Christian Müller, Marcel Schmiel, Nazanin Alavinejad, Agnieszka Rumińska, Li Zhang, Clara Ludwig, Lukas Maas, Miloš Nikolić, Lucia A. Torres Fernández, Graziella Bosco, Debora Puhleva, Amin Mokhlesi, Oliver Siefer, Felix John, Katja Höpker, Jan-Phillip Weber, Balazs Hegedüs, Fabian Doerr, Servet Bölükbas, Jens Peter Panse, Lars Hagmeyer, Cleidson Padua Alves, Lisa Werr, Maximilian Germer, Anna-Lena Schumacher, Magdalena Bogus, Cornelius Courts, Alexander Quaas, Reinhard Büttner, Giannicola Genovese, Maria Cartolano, Kjong-Van Lehmann, Martin Peifer, Roman K Thomas, Julie George

## Abstract

Platinum-based chemotherapy is the standard first-line treatment for patients with small cell lung cancer (SCLC), yet the scarcity of tumor specimens limits our understanding of how genetic and transcriptional heterogeneity interact to shape therapeutic resistance. To reconstruct tumor evolution *in vivo*, we leveraged a clinically annotated cohort of 85 SCLC patient-derived xenotransplant (PDX) models, and longitudinally profiled 33 models spanning distinct chemotherapy responses using mutation-based phylogenies and single-cell transcriptomics. Chemotherapy-sensitive tumors underwent extensive genomic evolution following treatment, driven by the selection of pre-existing subclones and the emergence of *de novo* clones harboring platinum-associated DNA damage. Prolonged chemotherapy exposure repeatedly selected alterations in SCLC-relevant genes, including *TP73* implicating these events in therapeutic resistance. Across models, genomic heterogeneity was linked to transcriptional diversification and adaptive changes in tumor cell states. Integrating single-cell analyses of evolutionary dynamics with optimal-transport modeling and experimental interrogation of cell-state dynamics during chemotherapy revealed convergent enrichment of epithelial–mesenchymal transition (EMT)-associated states across distinct models, linking the persistence of resistant clones with transcriptional adaptation. Together, these findings establish an experimental *in vivo* framework for modelling clinically relevant tumor evolution in SCLC and elucidate how therapy-driven clonal selection links genomic heterogeneity to transcriptional cell-state adaptation to shape SCLC chemotherapy resistance.

## Introduction

Small cell lung cancer (SCLC) is one of the most aggressive human malignancies, characterized by rapid growth, early metastasis, and a poor 5-year overall survival rate of less than 7%^1–6^ . First-line platinum–based chemotherapy is the standard treatment leading to high response rates but nearly all patients relapse within months^7^ . Recent advances in immunotherapy have conferred a clinical benefit, leading to the inclusion of anti-PD-L1 agents into first-line platinum–etoposide therapy^8,9^. Furthermore, DLL3-targeting T-cell engagers have shown remarkable clinical activity in relapsed SCLC, with emerging clinical trials supporting their potential in the first-line setting in combination with the current standard of platinum, etoposide, and PD-L1 inhibition^10,11^. Thus, despite these recent advances, platinum-based regimens remain the cornerstone of first-line therapy for patients with SCLC. This is due to a remarkable initial chemosensitivity – rarely observed in other cancers – that enables high response rates of around 60–70%, which is almost invariably followed by rapid development of therapy-refractory disease; a paradox that remains mechanistically incompletely understood, emphasizing the need to define the molecular basis of resistance to platinum-based chemotherapy^7,12^.

Previous studies led by us and others have provided important insights into the complex biology of SCLC revealing near-universal bi-allelic inactivation of the tumor suppressors *TP53* and *RB1*, recurrent amplifications of *MYC* family genes and frequent alterations in genes involved in cell cycle regulation and chromatin remodeling^13–17^. Transcriptional profiling further pointed to the role of key lineage-defining transcription factors (TFs) that shape neuroendocrine differentiation in SCLC, leading to studies interrogating subtype-specific therapeutic vulnerabilities^3,18–21^. Furthermore, first studies have provided insights into molecular pathways associated with chemotherapy resistance, implicating epigenetic reprogramming, loss of neuroendocrine identity, and activation of WNT transcriptional programs^22–27^. In this regard, we recently conducted multi-regional and longitudinal genomic profiling of patients with SCLC to delineate tumor evolutionary trajectories under therapy, revealing a pseudo-clonal, highly homogeneous population at first diagnosis, and demonstrating how first-line chemotherapy selectively depletes dominant tumor clones, thereby unleashing ancestral lineages that drive clonal diversity at relapse, with alterations in *TP73* and *CREBBP*/*EP300* as key determinants of treatment response^13^.

Studies across diverse cancer types have shown that therapeutic resistance arises from the interplay between genomic heterogeneity and non-genetic transcriptional and epigenetic adaptations which drive clonal selection^28–31^. However, understanding how these processes are coordinated in space and time under clinically relevant chemotherapy remains challenging, because longitudinal patient-matched samples are rarely available, and limited tumor material obtained from patients with SCLC often precludes comprehensive molecular analyses. To enable deep molecular and functional profiling of SCLC patient tumors, we and others have therefore established patient-derived xenograft (PDX) models from tumor tissue and circulating tumor cells (CTCs), which preserve genomic and molecular features of the original tumors^32–34^. Here, we leveraged a clinically annotated in-house cohort of 85 SCLC xenotransplant models spanning diverse treatment histories to systematically investigate treatment-driven tumor evolution. By integrating phylogenies based on single nucleotide variants (SNVs) with single-cell transcriptomics, we aimed at tracking chemotherapy-associated clonal evolution capturing both genomic as well as transcriptomic phenotypes under chemotherapy. This multi-layered framework advances our previous study of genomic evolution in patient tumors^13^ by moving from clinical observation to experimentally controlled *in vivo* models, linking clonal dynamics to treatment-induced cellular adaptations under therapeutic pressure.

## Results

### Chemotherapy response is recapitulated across genomically and molecularly diverse SCLC xenotransplant models

To investigate treatment-driven evolutionary dynamics of SCLC tumors *in vivo*, we analyzed our clinically annotated in-house cohort of 85 PDX models for comprehensive molecular characterization. This cohort, established from either patient-derived tumor tissue (n=32) or circulating tumor cells (CTCs, n=53), predominantly comprised treatment-naïve tumors (n=62, 73%), as well as models obtained during therapy (n=2, 2%) and at relapse after first-line platinum-based chemotherapy (n=21, 25%, **Fig. 1a**, **Supplementary Table 1–2**). Genomic profiling confirmed for all SCLC models near universal loss of *TP53* and *RB1*, along with recurrent alterations in *TP73* (21.2%), *CREBBP*/*EP300* (30.6%), *KMT2D* (37.6%), *NOTCH1–4* (34.1%) and amplification of *MYC* family genes (18.8%), confirming earlier studies on larger patient cohorts^13,15^ (**Supplementary Table 3**, **Suppl. Fig. 1a**). Furthermore, consistent with previous reports^35,36^, dimensionality reduction analysis of transcriptomic profiles revealed segregation of tumor samples according to the predominant expression of neuroendocrine lineage-defining TFs (**Fig. 1b**). However, neither transcriptional nor genomic profiles of our models segregated by the clinical status at the time of sample collection (*treatment-naïve, during therapy, or at relapse*) or by the donor’s response to first-line chemotherapy, suggesting that baseline molecular phenotypes alone may not sufficiently explain interpatient differences in therapeutic outcome (**Suppl. Fig. 1b**).

**Fig. 1.**
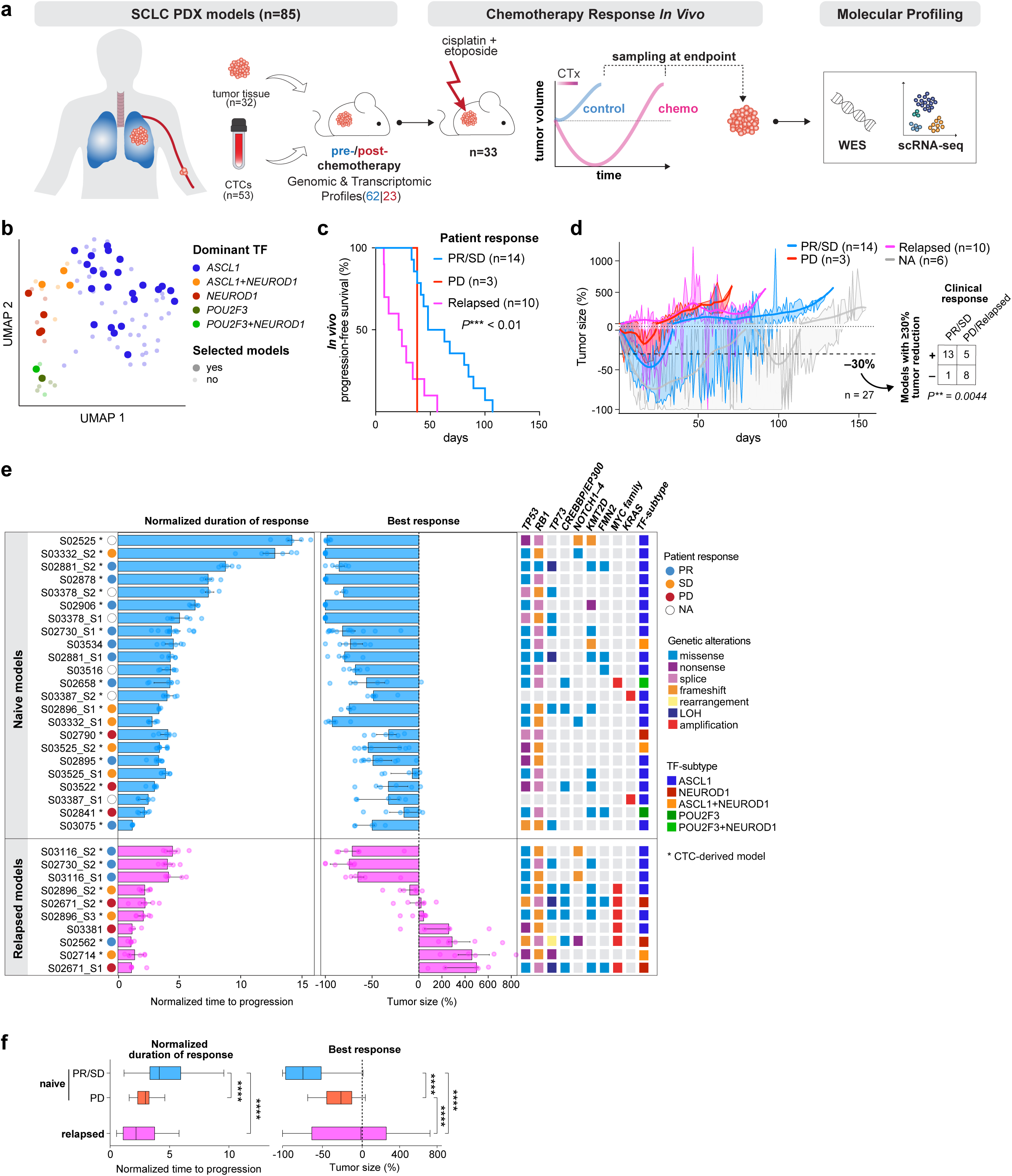
Molecular characteristics and chemotherapy response of SCLC xenotransplant models. **a**, Schematic overview of PDX cohort (n=85) established from pre- and post-treatment patient samples through engraftment of tumor tissue or circulating tumor cells (CTCs), with comprehensive genomic and transcriptional characterization of all models. *In vivo* chemotherapy responses assessed in 33 models, followed by in-depth molecular profiling with WES and single-cell RNA-seq of control and chemotherapy-treated recurrent tumors. **b,** UMAP embedding of bulk RNA-seq profiles from 85 PDX models colored according to the predominant expression of neuroendocrine lineage-defining transcription factors. Models selected for *in vivo* study are highlighted. **c–d,** *In vivo* functional assessment of chemotherapy responses in PDX models stratified by the clinical response of the corresponding patients: PR/SD (n=14, blue), PD (n=3, red), relapsed (n=10, magenta) and not available (NA, n=6, grey). (**c**) Kaplan–Meier analysis of progression-free survival (PFS) determined in models with available clinical response data (n=27). (**d**) Relative tumor growth across all models (n=33), categorized according to matched patient response. Contingency table showing models with ≥30% tumor shrinkage stratified by patient response (PR/SD vs PD/relapse). Fisher’s exact test, two-sided, *\*\*P* < 0.01. **e,** Responses to chemotherapy quantified for PDX models established from naïve and relapsed tumor samples displaying normalized duration of response (left) and best response (right), sorted by the composite score integrating both metrices (**Methods**). The clinical response of the corresponding patient is indicated by colored circles. CTC-derived models are marked with an asterisk (*); two models with LCNEC histology are indicated by #; 1 model established after the first cycle of chemotherapy is indicated by double dagger (^‡^). Key genetic alterations and dominant transcription factor (TF) subtypes for each model are annotated on the right. **f,** Box plots showing normalized duration of response (left) and best response (right), stratified by matched patient clinical response and by tumor sampling timepoint (treatment-naïve or at relapse). Pairwise Welch’s t-tests with Holm correction for multiple comparisons, \**P* < 0.05, \*\*\**P* < 0.001, \*\*\*\**P* < 0.0001.

To functionally characterize chemotherapy response *in vivo*, we selected 33 representative models derived from 23 patients capturing the broad spectrum of neuroendocrine lineage-defining TF expression patterns, genomic profiles, clinical responses to first-line platinum-based therapy, and sampling timepoints from both treatment-naïve tumors and relapsed disease following prior chemotherapy (**Fig. 1a–b, Suppl. Fig. 1a-b**). To model the platinum-etoposide regimen used in patients with SCLC, tumor-bearing mice were treated with etoposide and cisplatin for 2 weeks^37,38^. Tumor growth was monitored in both control- and chemotherapy-treated mice, and normalized to baseline tumor size at day 0, revealing marked variability in growth dynamics across this panel of models (**Suppl. Fig. 2, Methods**). Notably, this cohort included two treatment-naïve PDX models from a patient with LCNEC (S03387), featuring characteristics of a high-grade neuroendocrine tumor and *KRAS* amplifications, and revealing a comparably limited response to *in vivo* chemotherapy treatment. To further contextualize the *in vivo* chemotherapy response, we analyzed specifically SCLC PDX models with available clinical response data to first-line platinum-based therapy (partial response [PR], stable disease [SD], or progressive disease [PD]; n=27, here excluding both LCNEC models, **Supplementary Table 1**), revealing significant differences in tumor progression *in vivo* (Log-rank test *P*\*\*<0.01, **Fig. 1c**). Treatment-naïve PDXs derived from patients with clinical PR or SD showed increased chemotherapy sensitivity compared to those from patients with PD or those established at relapse, as evidenced by a higher proportion of models achieving ≥30% tumor shrinkage and lower area-under-the-curve (AUC) values, reflecting pronounced tumor regression (**Fig. 1d, Suppl. Fig. 3a, Supplementary Table 4–5)**. To further quantify these *in vivo* observations in analogy to clinical response metrics, we determined the duration of response (calculated as the time-to-progression/doubling time^34^, **Methods**), and the best response (defined as the minimum relative tumor volume; % of baseline) observed after completion of chemotherapy (day 14), with lower values indicating greater shrinkage (**Fig. 1e, Methods, Supplementary Table 5**). Both metrics demonstrated significantly enhanced responses and greater tumor shrinkage in models derived from patients with clinical benefit from platinum-based chemotherapy (**Fig. 1e–f**), confirming that our models faithfully recapitulate patient-specific therapeutic responses, consistent with previous studies of SCLC PDX models^32–34^. We further classified all 33 PDX models as “sensitive” (n =15), “intermediate” (n =5), or “resistant” (n=13) to chemotherapy based on a composite score integrating best response and duration of response (**Methods, Suppl. Fig. 2 and 3b, Supplementary Table 5**). Although models derived from relapsed patients showed reduced chemotherapy sensitivity, we note that *in vivo* responses varied considerably, with 3 relapsed models from patients with prior PR to first-line chemotherapy remaining responsive and classified as “sensitive”, which is in line with clinical observations that a subset of patients can benefit from platinum rechallenge^39–41^ (**Fig. 1e, Suppl. Fig. 3b**).

Altogether, our cohort enabled screening of 33 PDX models derived from 23 patients, demonstrating a strong concordance between *in vivo* response and corresponding clinical outcomes in the corresponding patients, thus corroborating previous findings from independent SCLC PDX cohorts^32,34^. We assessed whether key genomic alterations and transcription factor expression patterns across PDX models from all 23 patients were predictive of *in vivo* chemotherapy responses. However, no clear associations were observed for most frequent genomic alterations, (including ***KMT2D****, **NOTCH**, **CREBBP/EP300**, **TP73 and TP53***, **Suppl. Fig. 3c**). Notably, consistent with previous reports^13,22,23,25,26,42^, tumors with predominant *NEUROD1* (n=5) and *POU2F3* (n=2) expression, or *MYC* family amplifications (n=4) were associated with increased tumor progression *in vivo* – although these subgroups comprised only a small number of models (**Suppl. Fig. 3c**). These findings therefore suggest that baseline molecular features only partially explain therapeutic response, prompting us to investigate how tumors evolve under treatment and how chemotherapy-induced clonal dynamics contribute to treatment resistance and tumor recurrence.

### Chemotherapy-induced genomic evolution shapes transcriptional heterogeneity in SCLC

To elucidate how molecular features evolve under chemotherapy, we resolved genomic trajectories and transcriptional phenotypes at a single cell level, integrating these with *in vivo* response data. We collected for each chemotherapy-treated model typically two tumors at recurrence (chemo 1 and chemo 2, S2 and S3, respectively) along with one matched control tumor (S1), which were all subjected to high-coverage whole-exome sequencing (WES; n=95 samples) and single-cell RNA sequencing (scRNA-seq, n=96 samples, **Fig. 1a, Methods, Supplementary Table 2**).

WES data for all PDX tumors yielded a median coverage of 186× and a high tumor purity (median 99%, **Methods**) thus enabling detailed reconstruction of clonal dynamics (**Supplementary Table 6**). We determined cancer cell fractions (CCFs) for somatic mutations and assigned them to clonal populations using pairwise two-dimensional clustering (**Methods**), thereby reconstructing treatment-related phylogenetic trees for each model. C0 denotes the most recent common ancestral clone harboring mutations shared across all samples with CCFs at 100% (**Fig. 2a**), whereas subclones (C1–C2) were defined as clusters of mutations that were either sample-specific or present at CCFs below 100% (**Methods, Supplementary Table 6–7**). We thus identified both linear and branching evolutionary trajectories in a substantial proportion of our models (n=25/33 PDX models). When stratifying for *in vivo* treatment response, we observed that recurring tumors derived from chemotherapy-sensitive models exhibited more complex genome evolutionary processes involving both branching and linear evolution, resulting in significantly higher numbers of subclonal mutations and subclones, whereas resistant tumors showed limited clonal diversification (**Fig. 2b–c**). To further characterize these dynamics, we determined treatment-associated changes in CCFs for individual subclonal populations across all models and treated tumors (**Suppl. Fig. 3d**). Tumors were further classified by evidence of clonal selection, including *de novo* emergence of clones as well as the expansion or extinction of pre-existing subclones (**Fig. 2a**). Processes of clonal selection – particular *de novo* emergence of clones – were significantly enriched in chemotherapy-sensitive tumors compared to resistant models, with expansion of subclones beyond C1 becoming dominant in a subset of sensitive, but not in resistant tumors (Fisher’s exact tests, P<0.05, **Fig. 2d, Suppl. Fig. 4a**). In contrast, treatment of resistant models resulted in limited tumor evolution compared with sensitive models (**Fig. 2d**). Consistent with this observation, resistant or intermediate models exhibited a significantly lower number of subclonal mutations and subclones undergoing selection, defined as a substantial change in CCF of at least 30% (ΔCCF ≥ 30%) than chemotherapy-sensitive models (**Fig. 2e**). Notably, ancestral clones present prior to treatment (C0 or C1) accounted for a substantial fraction of the tumor cell population at relapse across all resistant and most sensitive models (**Suppl. Fig. 4a**), which is consistent with our prior observations from longitudinal analyses of tumor evolution in patients with SCLC^13^. Our data therefore demonstrate that *in vivo* tumor models not only recapitulate patient responses to chemotherapy, but also capture inherent subclonal genomic diversity and thus enable detailed investigation of evolutionary trajectories under treatment.

**Fig. 2.**
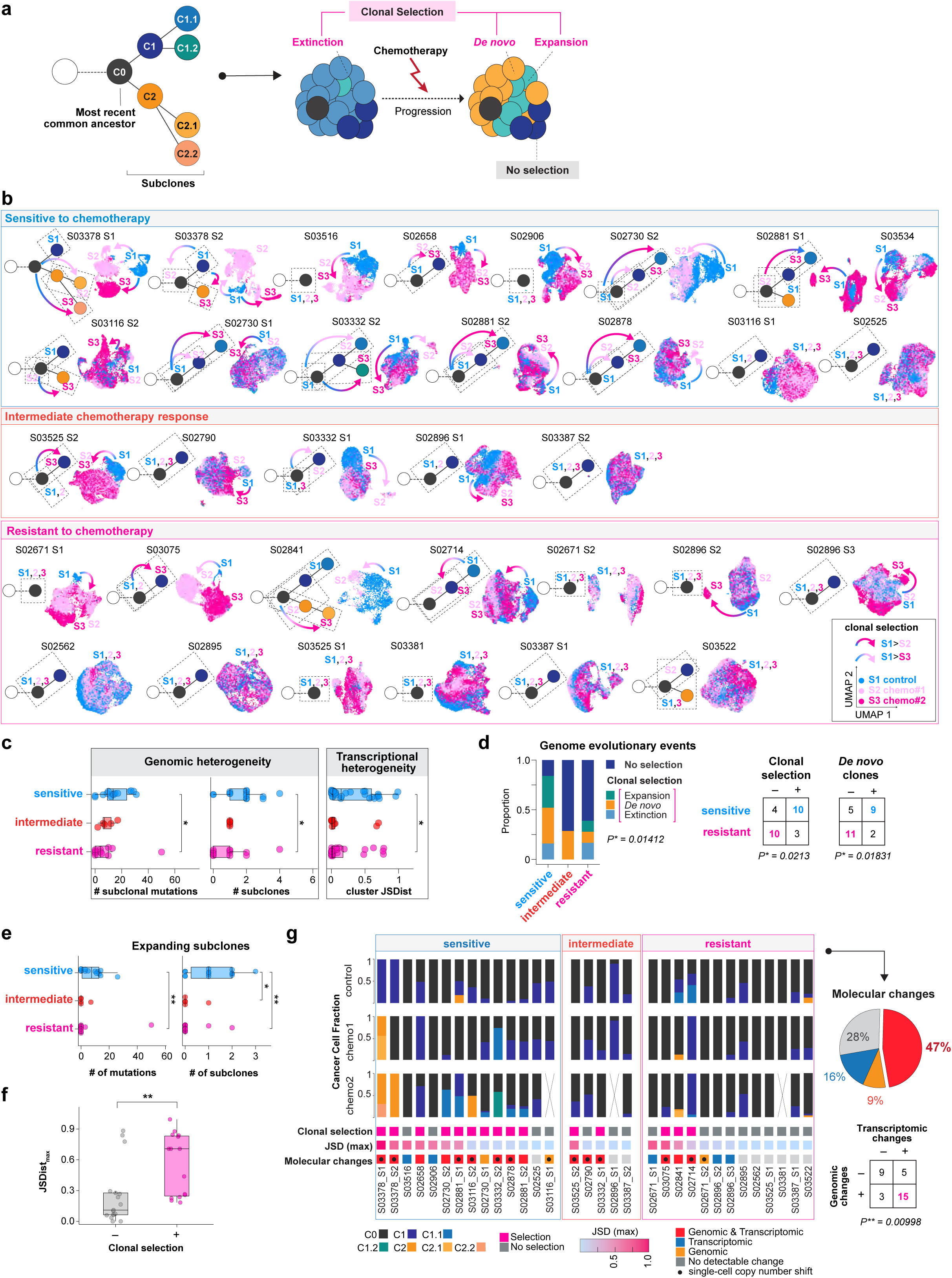
Integrated analysis of chemotherapy–driven genome evolution and transcriptional dynamics. **a**, Schematic illustration of mutation-defined clone phylogeny and clonal dynamics under chemotherapy depicting the most recent common ancestral clone (C0) and its descendant subclones (C1, C2, C3). Chemotherapy-associated clonal dynamics were categorized as either “no selection” or “selection” referring to the extinction or expansion of pre-existing clones and the *de novo* emergence of tumor clones. **b,** SNV-based phylogenetic trees displayed alongside single-cell transcriptomic UMAP embeddings for each PDX model, showing control (S1) and chemotherapy-treated samples (chemo 1 and 2; S2 and S3, respectively). Models are stratified according to their *in vivo* response to chemotherapy as sensitive, intermediate, or resistant. Phylogenetic trees are annotated as in (**a**). Dashed boxes indicate clones detected in the respective samples S1, S2, S3. Color-coded arrows indicate clonal evolutionary dynamics inferred from SNV-based and scRNA-seq data with colors scale representing the sampling timepoints S1 (blue), S2 (light pink) and S3 (pink). Models without arrows showed no evidence of clonal selection. **c,** Genomic and transcriptional heterogeneity across *in vivo* chemotherapy response groups. The number of subclonal mutations and SNV-defined subclones is quantified based on genomic data (left and middle panel). Jensen–Shannon distance (JSDist) between control and chemotherapy-treated tumors are shown for transcriptional data (right panel). Two-sided Wilcoxon rank-sum tests (for genomic data) and one-sided Wilcoxon rank-sum tests (for transcriptional data). \**P* < 0.05. **d,** Distribution of genomic evolutionary events across *in vivo* chemotherapy response groups (left). Contingency analyses comparing processes of clonal “selection” and *de novo* emergence of clones between sensitive and resistant models (right). Fisher’s exact tests; *P*\* < 0.05. **e,** Number of subclonal mutations (left) and subclones (right) undergoing selection (ΔCCF ≥ 30%) across sensitive, intermediate and resistant PDX models. Two-sided Wilcoxon rank-sum tests; \**P* < 0.05, \*\**P* < 0.01. **f,** Maximum Jensen–Shannon distance (JSDist) of transcriptional cluster compositions between control and chemotherapy-treated tumors, stratified by the presence or absence of genomic clonal selection (≥30% CCF shift or *de novo* subclones). Two-sided Wilcoxon rank-sum test; \*\**P*< 0.01. **g,** Integrated representation of genomic and transcriptional evolutionary processes across 32 models with available WES data following chemotherapy treatment. SNV-defined clonal composition of control (S1) and chemotherapy-treated samples (chemo 1 and 2; S2 and S3, respectively) is shown in the top panel, displaying the cancer cell fraction (CCF) for each sample. Crossed-out cases indicate missing biological replicates. SNV-based clonal dynamics (clonal selection) are annotated below. Transcriptional JSD is represented as continuous values, using the maximum JSD (JSDmax) across chemotherapy replicates for each model (see scale). Molecular changes summarizing genomic and transcriptomic heterogeneity are annotated, with additional indications of CN-based clonal dynamics inferred from single-cell RNA-seq (solid circles). Pie chart summarizes the distribution of PDX models exhibiting combined genomic and transcriptional evolution, transcriptional changes alone, genomic changes alone, or no detectable changes. The contingency table shows the association between genomic and transcriptional evolution. Fisher’s exact test, two-sided, \*\**P* < 0.01.

We next examined whether genome evolutionary processes impact transcriptional cell states. Single-cell RNA-seq yielded transcriptomic profiles from more than 360.000 cells derived from control- and chemotherapy-treated tumors across 33 PDX models, with an average of approximately 3,800 cells per tumor (median, 4217; IQR, 2220 – 5222) and a median of 3,620 detected genes (IQR: 3,088–4,146) and 11,170 UMIs (IQR: 8,817–14,414) following standard quality control filtering (**Methods, Supplementary Table 9**). To enable direct comparison of genomic evolution and transcriptional phenotypes, single-cell transcriptomes were visualized by UMAP alongside matched phylogenetic trees, with both analyses stratified by *in vivo* chemotherapy response (**Fig. 2b**). We first assessed intra-tumoral heterogeneity in each sample using pairwise Euclidean distances between cells, revealing higher levels in chemotherapy-treated tumors than in controls (**Suppl. Fig. 4b**). To further characterize treatment-associated changes in the tumor transcriptional landscape in each model, we quantified differences in transcriptional cluster composition between control and chemotherapy-treated samples using the Jensen–Shannon distance (JSDist, calculated as the square root of the Jensen–Shannon divergence (JSD), **Methods, Fig. 2c, Suppl. Fig. 4c**). Substantial levels of transcriptional heterogeneity determined by emergence of condition-specific clusters or JSD >0.1 were evident in 63.6% of all models (n=21/33, **Supplementary Table 10**). Stratification by treatment response revealed that, compared to resistant tumors, chemotherapy-sensitive models exhibited pronounced treatment-induced shifts in relative proportions of transcriptional clusters (**Fig. 2c**, **Suppl. Fig. 4c**, P<0.05).

Given the reported role of transcriptional plasticity in SCLC^22,23,26,27^, we examined whether chemotherapy alters lineage transcription factor (TF) states. Cells were assigned to key TF subtypes based on *ASCL1*, *NEUROD1*, *POU2F3*, and *YAP1* expression (**Methods**), revealing both single- and dual-positive states within tumors and substantial co-expression of key TFs in individual cells across all tumor models (**Suppl. Fig. 4d**). However, chemotherapy treatment did not alter lineage TF states, and TF-subtype distributions remained largely unchanged across control and treated tumors from all 33 models, despite increased transcriptional variability in chemotherapy-sensitive models (**Suppl. Fig. 3d–e**). Thus, processes beyond neuroendocrine lineage-defining TF expression likely contribute to treatment-induced transcriptional heterogeneity. Given the extensive genomic evolution observed following treatment, particularly in chemotherapy-sensitive models, we hypothesized that these evolutionary processes are key drivers of the observed transcriptional heterogeneity.

To complement the SNV-based phylogenies with genomic information at single-cell resolution we inferred copy number (CN) alterations from transcriptional profiles of each cell, enabling direct mapping of genomic changes onto transcriptional clusters (**Methods, Supplementary Table 11**). We thus identified novel CN-defined clusters, which were absent or rare in controls (≤1% of cells), but emerged following chemotherapy treatment in 6 of 15 sensitive and 2 of 5 intermediate models, but in none of the 13 resistant models, indicating a significant association with chemotherapy sensitivity (Fisher’s exact test, P<0.05; **Suppl. Fig. 4f**). Furthermore, all eight models with novel CN-defined clusters also exhibited SNV-based clonal selection, with a significant association between the emergence of novel CN-defined clusters and SNV-based evidence of clonal evolution (Fisher’s exact test, P<0.001; **Suppl. Fig. 4f**). This concordance between CN- and mutation-based evolutionary processes potentially reflects their biological interdependence, and support the use of transcriptome-inferred CN alterations as an additional layer for tracking genome evolution at single-cell resolution. We next investigated the relationship between genomic evolution and treatment-associated transcriptional changes across models with matched WES and scRNA-seq data (n=32, **Methods, Supplementary Table 2**). Models exhibiting SNV-based clonal selection showed significantly greater transcriptional divergence, as measured by the maximum JSDist (**Fig. 2f**). Integrating SNV- and CN-based evolution with transcriptional changes assessed for chemotherapy-treated and control tumors revealed that nearly half of our models (n = 15/32, 47%), most of which exhibited sensitive or intermediate response to chemotherapy (9/14 sensitive, 3/5 intermediate, 3/13 resistant; Fishers exact test, P = 0.0359), exhibited both genomic evolution and transcriptional changes, whereas isolated genomic or transcriptional changes were less frequent (**Fig. 2g, Supplementary Table 12**). Genomic and transcriptional changes were significantly associated across models (Fisher’s exact test, P < 0.01; **Fig. 2g**), demonstrating frequent co-occurrence and indicating a tight coupling between both processes. Thus, increased levels of transcriptional heterogeneity identified following chemotherapy treatment were predominantly accompanied by underlying genomic alterations, supporting a direct link between genome evolution and transcriptional diversity.

Collectively, we demonstrate that *in vivo* models for SCLC enable high-resolution dissection of tumor evolution under therapeutic pressure. Across a diverse panel of PDX models recapitulating various clinical responses to chemotherapy, we identified that chemotherapy-sensitive tumor models undergo extensive genomic evolution following platinum-based chemotherapy, including both selection of pre-existing subclones and the emergence of *de novo* clones. These genome evolutionary processes were accompanied by pronounced transcriptional reprogramming, but did not induce changes in neuroendocrine cell fate following platinum-based chemotherapy. Instead, underlying genomic heterogeneity emerged as a central driver of broader transcriptional diversity and subsequent cell state adaptations after treatment.

### Chemotherapy Drives Mutational Processes Associated with Recurrent SCLC Alterations

To further delineate how genomic alterations shape tumor evolutionary trajectories, we examined the spectrum of mutations arising in response to chemotherapy (**Supplementary Table 7**). We first analyzed single base substitution (SBS)^43^ to dissect mutational processes following treatment (**Supplementary Table 13, Methods)**. Consistent with previous reports^13,44–46^, clonal mutations were dominated by tobacco-associated processes (SBS4, SBS29, SBS92), endogenous ageing signatures (SBS1, SBS5), contributions from DNA repair defects (SBS3, SBS36) and APOBEC activity (SBS2, SBS13). By contrast, subclonal mutations – together with persistent ageing signatures and increased APOBEC activity^47^ – were enriched for platinum-associated signatures (SBS31, SBS35); particularly in models from previously treated patients, reflecting mutagenic imprints of prior chemotherapy exposure (**Fig. 3a, Suppl. Fig. 5a**). Notably, in 5 models these platinum footprints also emerged *de novo* after 2 cycles of *in vivo* treatment (**Suppl. Fig. 5a**), highlighting the potential of our models to capture chemotherapy-induced mutations. Together, these data show that chemotherapy induces platinum-associated DNA damage and activates additional processes such as APOBEC activity, both of which drive genomic evolution under treatment and promote tumor progression in our *in vivo* models.

**Fig. 3.**
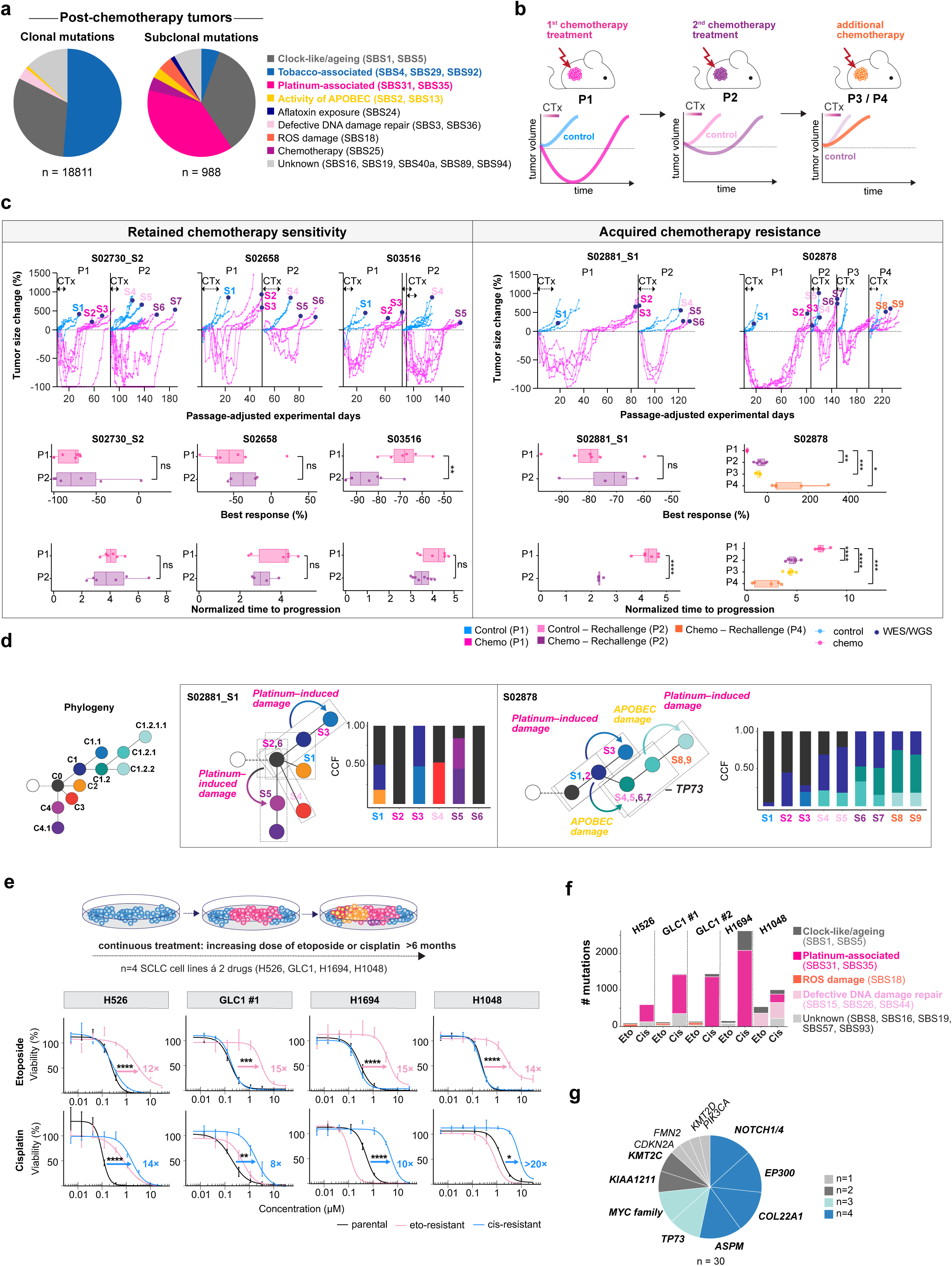
Mutational processes and recurrent SCLC alterations associated with prolonged chemotherapy treatment. **a**, Pie chart showing the relative contributions of mutational signatures based on clonal and subclonal mutations pooled separately across 32 PDX models, considering only mutations detected in post-chemotherapy tumor samples. Clonality was determined across all available samples per patient for phylogenetic reconstruction. **b,** *In vivo* cisplatin/etoposide rechallenge in five chemotherapy-sensitive PDX models. Schematic of the experimental design for two cylces of *in vivo* chemotherapy (CTx) followed by tumor impantation acrosse subsequent passages (up to P4, upper panel). Tumor growth, best response and normalized time to progression across passages are shown for models retaining chemotherapy sensitivity (S02730_S2, S02658 and S03516, left panel) and those acquiring resistance (S02881_S1 and S02878, right panel). For resistant models, SNV-based phylogenetic trees illustrate longitudinal clonal evolution and mutational signatures. CCF plots describe tumor-specific clonal compositions. Pairwise Welch’s t-tests with Bonferroni correction for multiple comparisons for S02878, \**P* < 0.05, \*\**P* < 0.01, \*\*\**P* < 0.001, \*\*\*\**P* < 0.0001. **c,** *In vitro* dose escalation in SCLC cell lines. Schematic of continuous exposure to increasing cisplatin or etoposide concentrations over >6 months (upper panel). CTG viability dose-response curves for parental and resistant cell line derivatives (H526, GLC1 #1, H1694 and H1048) following cisplatin (cis) or etoposide (eto) treatment. Data represent mean ± SEM from at least three biological replicates per condition; curves fitted using four-parameter logistic regression. Arrows indicate increased IC_50_ values in resistant derivatives with corresponding fold changes indicated numerically, two-sided Welch’s t-tests on log_10_-transformed IC_50_ values (**Suppl. Fig. 3e**), \**P* < 0.05, \*\**P* < 0.01, \*\*\**P* < 0.001, \*\*\*\**P* < 0.0001. **d,** Stacked bar plots showing the number of *de novo* mutations acquired in cisplatin- and etoposide-resistant derivatives of SCLC cell lines (GLC1 #1, GLC1 #2, H1048, H1694, H526), categorized by their assigned mutational signatures. **e,** Pie chart showing the proportion of recurrent SCLC-specific gene alterations identified across n=10 resistant SCLC cell lines.

Although substantial numbers of mutations were acquired after two cycles of chemotherapy, particularly in sensitive models, no recurrently affected genes were found across all models. Instead, acquired somatic alterations converged on pathways involved in proliferation and stress adaptation indicating that *in vivo* chemotherapy treatment primarily selects for alterations in broader functional modules rather than individual genes (**Suppl. Fig. 5b**).

Since patients with SCLC typically receive 4–6 cycles of chemotherapy^48^ and are followed over longer clinical timescales than feasible in murine models, we next performed *in vivo* rechallenge experiments to interrogate how sustained chemotherapy exposure shapes clonal evolution. Five chemotherapy-sensitive models were harvested at recurrence following 2 cycles of chemotherapy (Passage 1, P1), re-implanted into naïve hosts, and randomized to either no further treatment or additional cisplatin/etoposide treatment, thus enabling a cumulative *in vivo* exposure of 4–8 cycles of chemotherapy (across passages P2–4) with longitudinal monitoring of tumor growth for up to 8 months (**Fig. 3b**). 3 of 5 models retained chemotherapy-sensitivity (S02730_S2, S02658 and S03516) over two *in vivo* passages and treatment cycles, consistent with the clinical observation that patients with SCLC can benefit from chemotherapy rechallenge^1,39–41,48^. By contrast, chemotherapy re-challenge in 2 models revealed signs of acquired resistance with significantly reduced time to progression (S02881_S1 and S02878) and depth of response (S02878) and (**Fig. 3b, Supplementary Table 3**). Longitudinal genome profiling revealed that models retaining chemotherapy sensitivity exhibited only limited clonal evolution with very few subclonal mutations emerging after P1 and minimal additional clonal changes after P2, whereas models that acquired chemotherapy resistance showed a substantial increase in subclonal mutation burden, accompanied by extensive clonal diversification and branching evolutionary architectures (**Fig. 3b**, **Suppl. Fig. 6a–b**). We examined genomic alterations across all tumors from these models, and identified in the resistant model S02878 clonal remodeling coinciding with focal copy number loss at the 1p36.32 locus encompassing *TP73*. Complementary WGS confirmed genomic rearrangements leading to *TP73* loss, and complementary single-cell copy number inference from scRNA-seq confirmed *TP73* deletion in all tumors after two *in vivo* treatment regimens, suggesting a potential role for *TP73* loss in driving chemotherapy resistance^49,50^ (**Fig. 3b, Suppl. Fig. 6c–d)**.

To further study how prolonged chemotherapy exposure selects for recurrent genomic alterations – and to overcome the cycle-limited nature of our *in vivo* studies – we conducted complementary *in vitro* studies in established SCLC cell lines. We exposed cisplatin- and etoposide-sensitive SCLC cell lines (GLC1, H1694, H1048, H526; GLC1 included as duplicate runs) to stepwise escalating doses of either cisplatin or etoposide, thus enabling continuous chemotherapy treatment over 6-8 months (**Fig. 3c, Suppl. Fig. 6e, Methods**). Prolonged drug exposure led to a robust induction of therapy resistance across all cell lines, with at least 5-fold increases in IC_50_ (**Fig. 3c**, **Suppl. Fig. 6e, Supplementary Table 14**). Notably, 3 cisplatin-treated cell lines also acquired cross-resistance to etoposide. WES analysis of all resistant cell lines revealed substantial accumulation of mutations, with hundreds to thousands *de novo* mutations emerging in cisplatin-resistant cell lines (**Fig. 3d**) which were predominantly shaped by platinum-associated signatures (SBS31, SBS35, **Supplementary Table 13)**. Mutational footprints of etoposide-exposed cell lines on the contrary revealed ROS (SBS18), DNA repair–defect (SBS15, SBS26, SBS44), and clock-like (SBS1, SBS5) processes, consistent with double-strand breaks and oxidative stress caused by topoisomerase II inhibition^51,52^ (**Fig. 3d**). Among acquired gene mutations, recurrent genetic alterations were identified affecting key genes described for SCLC^15^, including mutations in *EP300*, *NOTCH1*, *KMT2C*, *KMT2D*, *COL22A1*, *ASPM* and *FMN2*, as well as copy number alterations involving *TP73*, *MYC* and *MYCN* (**Fig. 3e, Supplementary Table 15**).

Collectively, our data demonstrate that two cycles of platinum-based chemotherapy drive clonal diversification and *de novo* mutagenesis *in vivo*, accompanied by the emergence of dominant tumor clones harboring extensive platinum-associated genomic scars. Prolonged chemotherapy exposure *in vivo* and *in vitro* further uncovered selection of key recurrent SCLC-relevant alterations, supporting their contribution to therapy resistance. Although short-term *in vivo* treatment did not reveal recurrent gene alterations across PDX models, platinum-damaged clones emerged as dominant populations, and platinum-associated mutations converged on broader functional pathways. These findings may therefore suggest that chemotherapy-driven genome evolution initially promotes transcriptional adaptations to treatment, which subsequently facilitate the acquisition and selection of resistance-associated genomic alterations.

### Chemotherapy-resistant clones with platinum-induced mutational signatures exhibit activated epithelial-to-mesenchymal transition (EMT) programs

We next sought to study the selective advantage conferred by dominant tumor clones harboring platinum-induced mutational signatures by characterizing their associated transcriptional programs. We integrated SNV-informed tumor evolution with CN clones inferred from scRNA-seq data to robustly map clonal architecture onto transcriptional states, demonstrating concordance between SNV-defined and CN-defined clones in three cases (**Fig. 4a, Suppl. Fig. 7a–b, Methods**). In the chemotherapy-resistant model S02714, platinum-associated mutational signatures were detected in a pre-existing subclone that expanded and became dominant under chemotherapeutic pressure, whereas in chemotherapy-sensitive models (S02881_S1 and S02878) platinum scars were induced *de novo* after *in vivo* chemotherapy treatment. In an additional fourth case (S02730), we further analyzed matched PDX models established from the same patient at first diagnosis and after chemotherapy, with platinum signatures exclusively detected in the relapse but not treatment-naïve model, which enabled the assignment of platinum-damaged subclones to specific transcriptional cell populations. Across these models, tumor evolution was associated with the enrichment or emergence of platinum signature-bearing clones and cell clusters under chemotherapy, linking these mutational processes to the development of resistance. Using gene set enrichment analysis of Hallmark gene sets^53^, we analyzed transcriptional clusters expanding at relapse that were assigned to platinum-based genomic scars (**Methods**). While individual models exhibited distinct transcriptional changes involving metabolic, inflammatory, and cell cycle programs, epithelial–mesenchymal transition (EMT) emerged as a recurrent feature, showing significant upregulation in 3 of 4 models (**Fig. 4a–b**; **Suppl. Fig. 7c**). This analysis may thus point to the relevance of EMT-associated cellular states in shaping tumor evolutionary trajectories and promoting clonal outgrowth, consistent with recent studies implicating EMT in this process^54^. We further interrogated this program at single-cell resolution applying both the Hallmark EMT and previously validated EMT gene signatures^55^ (**Methods**), and confirmed in two models that tumor cells associated with platinum-induced damage display increased EMT scores (**Fig. 4b, Suppl. Fig. 7b**). We further examined transcriptional changes in SCLC cell lines with acquired resistance to either etoposide or cisplatin following prolonged *in vitro* chemotherapy exposure (**Fig. 3c**). Across cell lines with acquired resistance, recurrent transcriptional alterations predominantly involved EMT and inflammatory signaling pathways (**Suppl Fig. 7d–e**), which were observed not only in cells with acquired platinum damage, but also in cell lines treated with etoposide, indicating that distinct chemotherapeutic agents can converge on similar transcriptional programs associated with acquired resistance.

**Fig. 4.**
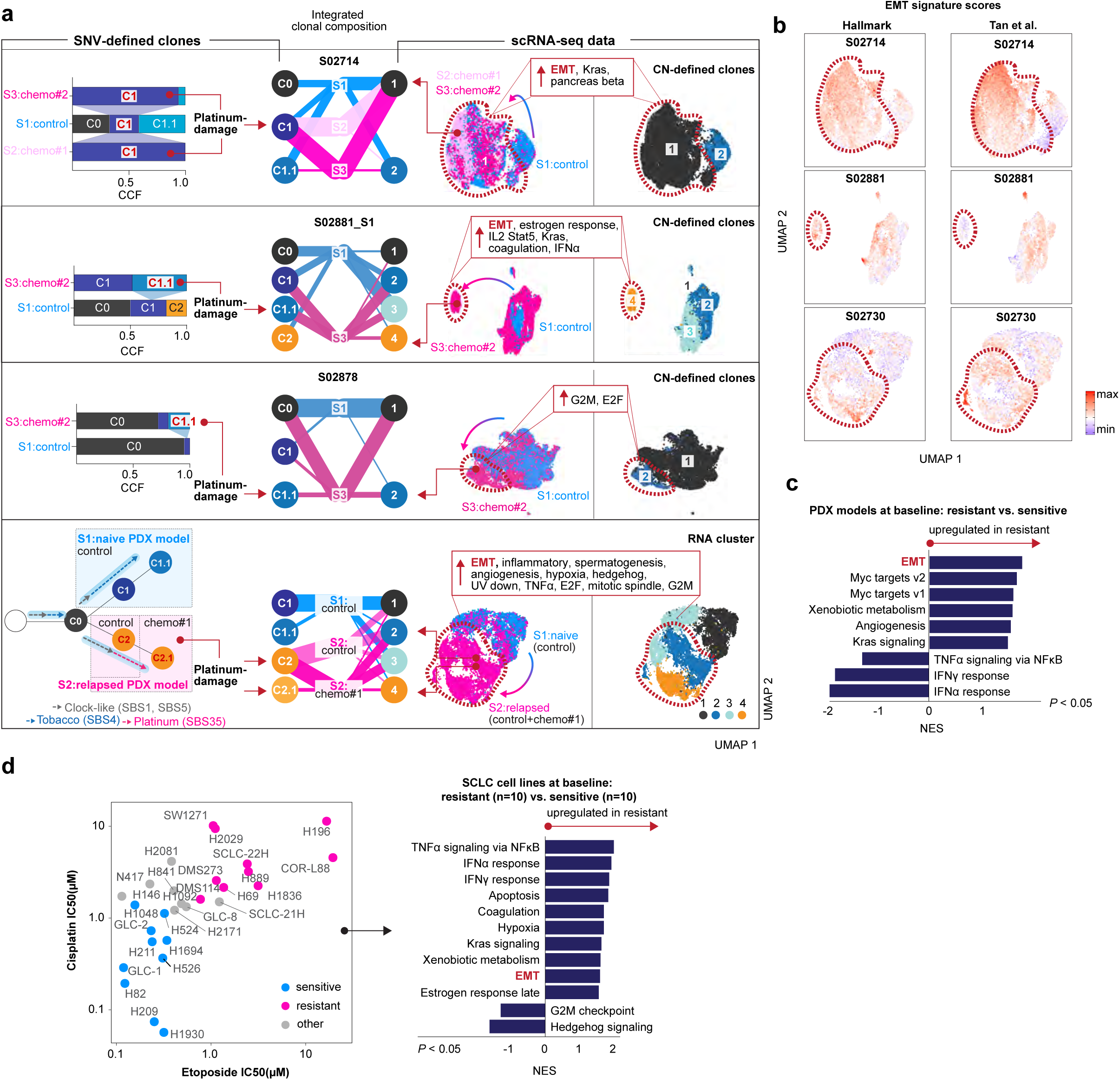
Transcriptional programs associated platinum-induced evolution and chemotherapy resistance. **a**, Integration of SNV-defined clonal evolution with single-cell transcriptomic states across four SCLC PDX cases (S02714, S02881_S1, S02878 and S02730). SNV-defined clonal compositions and phylogenies illustrate expanding tumor clones harboring platinum-associated mutational damage (left). Mapping of SNV-defined clones to single-cell RNA-seq data is based on their correspondence with scRNA-seq-inferred copy-number (CN) clusters or transcriptional clusters. Alluvial plots show the relative proportions of SNV-defined clones and corresponding CN clusters (S02714, S02881_S1 and S02878) or transcriptional clusters (Leiden; S02730) across samples (middle). UMAP representations show transcriptional states colored by sample identity or CN/RNA cluster assignment (right). Platinum-associated clones are indicated by red arrows, and their corresponding cell populations are outlined in red. Significantly enriched (*P* < 0.05) transcriptional programs associated with mapped platinum-damaged populations are indicated. **b,** EMT signature scores (Hallmark EMT and Tan et al.^55^) projected onto the corresponding single-cell UMAP embeddings for S02714, S02881_S1 and S02730. Red dashed outlines indicate cell populations associated with platinum-damaged clones. **c,** Gene set enrichment analysis (GSEA) of baseline bulk RNA-seq profiles comparing chemotherapy-resistant and chemotherapy-sensitive PDX models. Significantly enriched Hallmark pathways are ranked by normalized enrichment score (NES; *P* < 0.05). **d,** Cisplatin and etoposide IC_50_ values plotted against each other for 28 SCLC cell lines, with dual-sensitive and dual-resistant groups indicated (left) defined using drug-specific median IC_50_ values. GSEA comparing baseline transcriptomic profiles of intrinsically chemotherapy-resistant (n = 10) and chemotherapy-sensitive (n = 10) cell lines (right). Significantly enriched Hallmark pathways are ranked by NES (*P* < 0.05).

While platinum-associated genomic scars enabled us to track associated transcriptional states of outgrowing tumor clones, we next sought to investigate whether – irrespective of platinum-induced genomic damage – similar cellular states emerge across other model systems. Indeed, Gene Set Enrichment Analysis (GSEA) of baseline transcriptomic profiles from resistant versus sensitive SCLC PDX models identified EMT-related programs as the most strongly enriched pathways in resistant models, followed by *MYC*-related programs based on normalized enrichment scores (NES, **Fig. 4c, Suppl. Fig. 7f**). To further investigate whether these steady-state transcriptional programs extend beyond PDX models, we additionally profiled a panel of 28 SCLC cell lines and assessed their *in vitro* sensitivities to chemotherapy (**Supplementary Table 14**). Across cell lines expressing diverse neuroendocrine lineage transcription factors, we observed a broad range of sensitivities to cisplatin and etoposide, allowing classification into dual-sensitivity or dual-resistance groups (**Methods, Fig. 4d, Suppl. Fig. 7g**). Consistent with the findings in PDX models, baseline transcriptomic profiles of chemotherapy-resistant SCLC cell lines showed enrichment of EMT-associated transcriptional programs (**Fig. 4d**).

Altogether, our analyses reveal how genome evolutionary processes are accompanied by transcriptional adaptations which recurrently point to activation of EMT programs not only in platinum-damaged clones, but across the majority of chemotherapy-resistant SCLC models and SCLC cell lines. These findings suggest that, beyond specific genomic alterations, EMT-associated programs provide a permissive cellular state that promotes tumor adaptation and progression under therapeutic pressure, consistent with recent studies highlighting the role of EMT in driving tumor heterogeneity^54^.

### Cell-state trajectories and adaptive programs associated with chemotherapy resistance

To obtain a broader understanding of cellular adaptations associated with chemotherapy sensitivity and resistance, we next interrogated how within treatment response groups single tumor cells evolve under chemotherapeutic pressure. Joint embedding of all single-cell transcriptomes revealed clustering primarily by patient rather than treatment condition, indicating that inter-patient variability masks shared therapy-induced transcriptional patterns (**Fig. 5a**). We therefore investigated for each model the emergence of novel cellular transcriptional states among chemotherapy-treated tumor samples using k-nearest neighbor analysis (**Methods**), revealing that the distribution of newly emerging cellular states was shifted towards higher proportions in chemotherapy-sensitive tumors following treatment (One-sided Kolmogorov-Smirnov test, P* = 0.045, **Supplementary Table 10**). Using a threshold of >15% novel cells, novel cellular states were significantly more prevalent in chemotherapy-sensitive than chemotherapy-resistant models (Fisher’s Exact Test, P<0.05. **Fig. 5b, Methods**). Differentially expressed genes in newly emerging clusters from chemotherapy-sensitive tumors were found to be mostly enriched for adaptive transcriptional programs related to EMT/migration/extracellular matrix (ECM)/tissue integrity, metabolism/hypoxia and cell death/p53/stress. Chemotherapy-resistant tumors showed enrichment mostly in cell cycle/proliferation (**Suppl. Fig. 8a–b, Methods, Supplementary Table 16**). Notably, several of these functional programs, particularly those related to EMT/migration/ECM/tissue integrity, showed increased activity after chemotherapy in specific subpopulations of sensitive models, while related states were already detectable at baseline in subsets of chemotherapy-resistant models (**Methods**, **Suppl. Fig. 8b–d**), supporting the notion that resistant tumors at steady-state already feature transcriptional states primed for chemotherapy resistance (**Suppl. Fig. 8e, Suppl. Fig. 7f**).

**Fig. 5.**
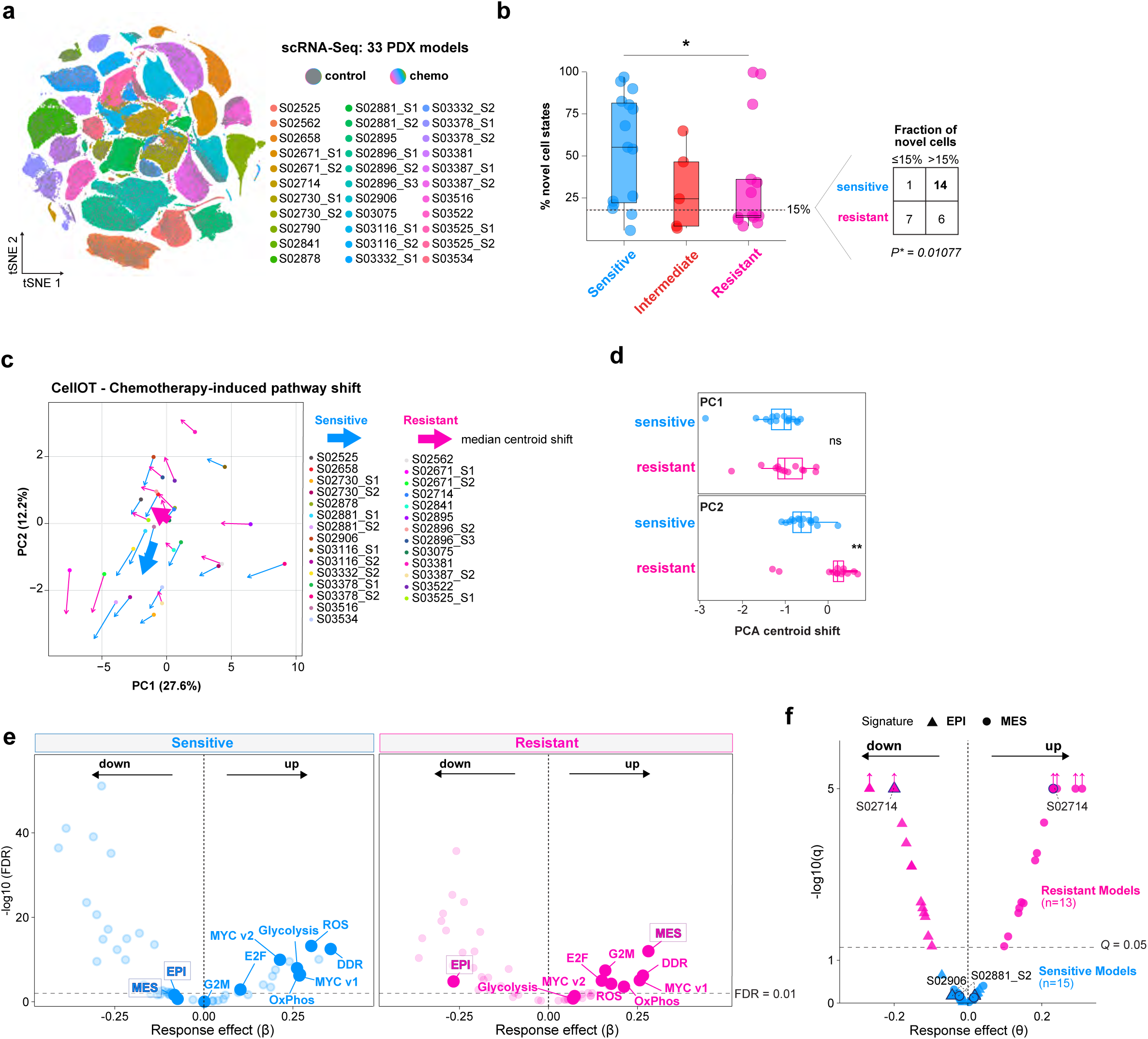
Modeling chemotherapy-induced cell-state transitions reveals response-specific adaptation in SCLC. **a**, Joint tSNE embedding of single-cell transcriptomes from control- and chemotherapy-treated tumors across 33 PDX models, colored by PDX model identity and treatment condition. **b,** Percentage of novel transcriptional cell states in chemotherapy-treated tumors across sensitive, intermediate and resistant PDX models, defined based on their local 30-nearest-neighbor (kNN) relationships relative to control cells. Statistical significance was assessed using a one-sided Kolmogorov–Smirnov test, \**P* < 0.05. Dashed line indicates the 15% threshold used for descriptive classification. The contingency table compares the frequency of models with >15% novel cells between chemotherapy-sensitive and resistant groups; two-sided Fisher’s exact test, \**P* <0.05. **c,** Principal component analysis (PCA) of CellOT-inferred chemotherapy-induced changes in Hallmark pathway activity and validated EMT signatures. Arrows connect control and CellOT-predicted chemotherapy-treated centroids for individual models; thick arrows indicate median centroid shifts within each response group. **d,** Model-level centroid shifts along PC1 and PC2 for chemotherapy-sensitive and -resistant PDX models. Shifts represent differences between the median coordinates of control and CellOT-predicted chemotherapy-treated cells. Each dot represents one model. Group differences were assessed using two-sided Wilcoxon rank-sum tests, \*\**P* <0.01 **e,** Response group-specific CellOT-inferred chemotherapy-induced pathway shifts in sensitive (left) and resistant (right) models, based on Hallmark and validated EMT signatures. Volcano plots show median pathway effect estimates (β) and corresponding FDR-adjusted significance; dashed lines indicate FDR = 0.01. **f,** Model-specific CellOT effect estimates (θ) for mesenchymal (MES) and epithelial (EPI) transcriptional programs in chemotherapy-sensitive and -resistant models. Arrows indicate models with −log_10_(q) values exceeding the plotting cut-off; these values are capped at 5. PDX models selected for longitudinal analyses (Fig. 6) are indicated. Dashed line indicates *Q* = 0.01.

To further disentangle treatment-driven state transitions from patient-specific variation, we applied a perturbation modeling framework based on optimal transport, using CellOT^56^ (**Methods**). As both control and chemotherapy-treated tumor samples represent static snapshots of dynamically evolving cell states under therapeutic pressure, this approach enabled us to infer chemotherapy-associated state transitions for each single cell. CellOT models were trained specifically on PDX models exhibiting pronounced sensitivity or resistance to chemotherapy, excluding xenotransplants with intermediate responses (**Suppl. Fig. 3a**). We first trained CellOT using a feature set comprising gene set variation analysis (GSVA)-derived Hallmark pathway scores^53^ and the validated EMT gene signatures from Tan et al.^55^, thereby capturing both broad transcriptional adaptations and specific EMT-associated states. We calculated PDX-specific centroid shifts from the control state to the CellOT-predicted chemotherapy-treated state and projected the directionality of treatment-associated cellular adaptations into a shared PCA space (**Fig. 5c, Supplementary Table 17**). We thus observed for chemotherapy-sensitive and -resistant PDX models pronounced transcriptional shifts, with largely consistent directions among models within each response group (**Fig. 5c**). These patterns were further characterized by quantifying shift magnitude and coordination (**Suppl. Fig. 8f, Methods**). The direction of transcriptional shifts differed significantly between response groups, primarily along PC2, with sensitive models shifting toward negative and resistant models toward positive PC2 values (two-sided Wilcoxon rank-sum test, P = 0.0043; **Fig. 5d**). Positive PC2 loadings were predominantly associated with mesenchymal and cell cycle-related signatures, whereas negative loadings were marked by metabolic programs, suggesting that these transcriptional programs potentially contribute to the distinct response group-specific adaptation patterns (**Suppl. Fig. 8g**).

We next investigated shared and response-specific transcriptional adaptations across our PDX models, as inferred by CellOT (**Fig. 5e, Suppl Fig. 8h, Supplementary Table 18**). Consistent with the pathways identified earlier (**Suppl. Fig. 8a–b**), chemotherapy elicited a broadly shared transcriptional response across tumors, characterized by increased MYC target, DNA damage response, and E2F signaling. However, the magnitude and direction of specific pathway shifts differed between response groups, with increased reactive oxygen species (ROS) and glycolysis programs in chemotherapy-sensitive models, whereas chemotherapy-resistant models exhibited increased G2M and mesenchymal programs (**Fig. 5e**, **Suppl. Fig. 8g**). To further resolve EMT-associated cellular state transitions, we applied a separate CellOT model trained exclusively on the Tan et al^55^. EMT gene signatures^55^. This focused analysis revealed a pronounced shift toward mesenchymal and away from epithelial transcriptional states in chemotherapy-resistant models, further supporting a role of EMT-related adaptations and acquired chemotherapy resistance (**Fig. 5f**).

Together, our data demonstrate that chemotherapy induces extensive transcriptional remodeling and the emergence of novel cell states in recurring tumors with initial sensitivity to chemotherapy. By contrast, resistant tumors largely retained pre-existing transcriptional programs. In this context, EMT-related transcriptional activity emerged consistently as a conserved feature of chemotherapy resistance/persistence with tumor clones expanding after chemotherapy enriched for EMT-associated transcriptional programs. This was evidenced from static transcriptomic snapshots of treatment-naïve and recurrent tumors after chemotherapy treatment, and was further confirmed through the computational framework of CellOT inferring cell state transitions associated with tumor evolution. Overall, EMT-related processes appear to be decisive controlling chemotherapy sensitivity and resistance.

### Longitudinal transcriptional dynamics under chemotherapy

Given the recurrent association of EMT-related programs with chemotherapy-induced cell state transitions, we next sought to experimentally investigate the temporal dynamics of tumor cell adaptation under ongoing chemotherapeutic pressure. To this end, we focused on three PDX models exhibiting distinct patterns of tumor evolution following *in vivo* chemotherapy treatment: one chemotherapy-resistant model (S02714), characterized by the expansion of pre-existing tumor clones harboring platinum-associated mutational signatures (**Fig. 4a**) and a pronounced EMT phenotype in recurrent tumors; and two chemotherapy-sensitive models with less pronounced EMT features, of which S02881_S2 exhibited both genome evolutionary processes and transcriptional changes, whereas S02906 showed transcriptional changes in the absence of apparent genomic evolution (**Fig. 2b and g, Suppl. Fig. 4f**, **Suppl. Fig. 9a**). Consistently, our CellOT model predicted an increased shift toward mesenchymal states following chemotherapy treatment in the resistant model, whereas this shift was significantly reduced in both sensitive models (**Fig. 5f**).

To further investigate how tumor cell states evolve in response to therapeutic pressure, we performed longitudinal single-nucleus RNA sequencing (snRNA-seq) of flash-frozen tumor tissues collected at multiple timepoints throughout treatment, from therapy initiation to tumor recurrence. Sampling was performed during the 2-week *in vivo* chemotherapy regimen (days 3 and 10, following completion of cycles 1 and 2, respectively), one week after completion of treatment (day 17), at the onset of tumor progression, at relapse (defined as regrowth to the initial tumor volume), and at the experimental endpoint (**Fig. 6a, Methods, Supplementary Table 4**). Although single-nuclei isolation from flash-frozen tumors yielded fewer nuclei for some samples particularly during treatment (median, 3,246; IQR, 1,268–4,945) and and snRNA-seq showed, as expected, lower transcriptomic coverage (median, 952; IQR, 638–1,296; and 1,313 UMIs; IQR, 842–1,908) compared with single-cell transcriptomic profiling of viable cells (**Methods, Supplementary Table 9**), recovered nuclei provided sufficient coverage for longitudinal studies of tumor cell states and their adaptations to chemotherapeutic pressure.

**Fig. 6.**
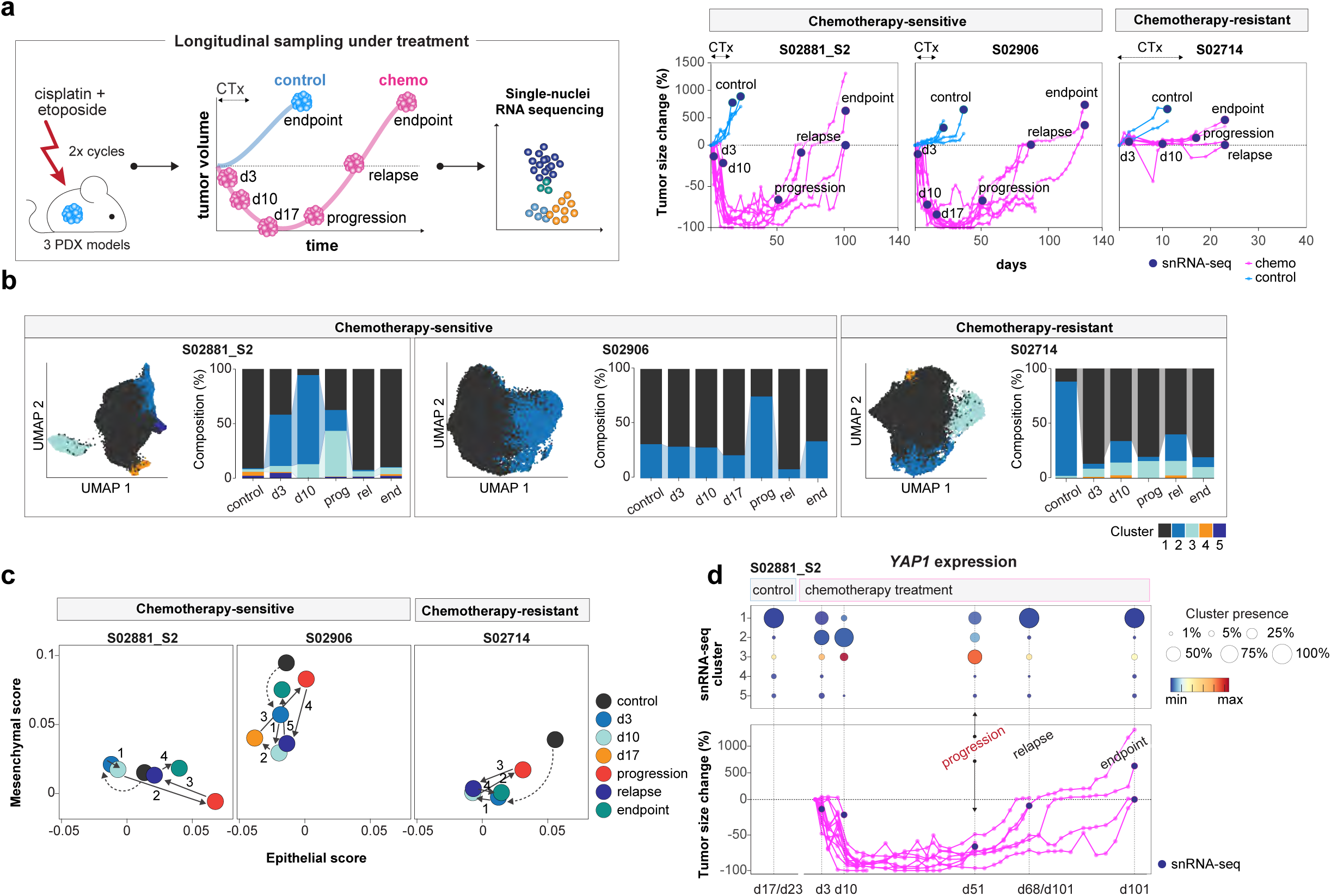
Longitudinal profiling reveals dynamic cell-state adaptation during chemotherapy and tumor regrowth. **a**, Schematic of longitudinal tumor sampling during chemotherapy (CTx) treatment. PDX models received two cycles of cisplatin and etoposide, with tumors collected during treatment (days 3 and 10), after treatment (day 17), at progression, at relapse, and at the experimental endpoint (left panel; **Methods**). Tumor growth curves for the chemotherapy-sensitive models S02881_S2 and S02906 and the chemotherapy-resistant model S02714 are shown on the right. Timepoints of tumor collection for snRNA-seq are indicated. **b,** UMAP embeddings for S02881_S2, S02906 and S02714, together with the transcriptional cluster composition across longitudinal sampling timepoints. Shaded transitions highlight the expansion of specific transcriptional clusters over time. Abbreviations: d, day; prog, progression; rel, relapse; end, endpoint. **c,** Longitudinal epithelial and mesenchymal transcriptional programs. Median epithelial and mesenchymal scores, based on Tan et al.^55^, were plotted across sampling timepoints to visualize shifts in epithelial/mesenchymal states during treatment and tumor regrowth. Numbers denote sequential transitions between tumor collection timepoints, as indicated by arrows. Dashed arrows indicate transitions in control tumors from baseline to the first collection timepoint during chemotherapy treatment. **d,** Longitudinal analysis of model S02881_S2 illustrating the transient emergence of *YAP1*-expressing cell states, highlighting the time of tumor progression. Circle sizes indicate the relative abundance of individual transcriptional clusters at each sampling timepoint, with colors representing *YAP1* expression levels (top). Corresponding tumor growth curve is shown below.

We first examined cell cycle distributions across treatment timepoints. Consistent with the nearly complete (100%) tumor regression observed in both chemotherapy-sensitive models (**Fig. 6a**), transcriptomic profiles of cell cycle states revealed a pronounced increase in the proportion of G1-phase cells after completion of two chemotherapy cycles (days 10–17), consistent with treatment-induced cell cycle arrest. In contrast, the chemotherapy-resistant model showed no marked change in cell cycle phase distribution, in line with the absence of tumor shrinkage and continued tumor cell proliferation after treatment (**Fig. 6a, Suppl. Fig. 9b**).

We next performed unsupervised clustering on transcriptional profiles across all timepoints within each PDX model to identify distinct tumor cell populations and track their dynamics throughout treatment. Consistent with our genomic findings (**Fig. 4a**), the chemotherapy-resistant model S02714 showed the expansion of a pre-existing transcriptional cluster upon treatment initiation, which predominated throughout treatment and persisted at the end of the treatment course (cluster 1, **Fig. 6b**). In contrast, both chemotherapy-sensitive models exhibited pronounced but transient transcriptional adaptations, characterized by the emergence, expansion and subsequent regression of timepoint- and response-dependent cell clusters (clusters 2 and 3 in S02881_S2, and cluster 2 in S02906). Accordingly, recurrent tumors in both chemotherapy-sensitive models largely recapitulated the transcriptional cluster proportions observed in baseline control samples, suggesting a re-establishment of a similar tumor cell state composition (**Fig. 6b**). Notably, in both models, transient cluster expansions were observed at the time of tumor progression, reflecting the timepoint when tumor growth resumed more than 40 days after treatment discontinuation following near-complete chemotherapy-induced tumor regression.

Given the EMT-associated cell state transitions identified above, we assessed the distribution of EMT-related transcriptional programs across the annotated tumor cell clusters. Despite distinct baseline epithelial (E) and mesenchymal (M) profiles, with S02906 displaying more mesenchymal and S02881_S2 more epithelial features, both chemotherapy-sensitive models exhibited a transient upregulation of epithelial features in expanding cell clusters at the onset of tumor progression (**Fig. 6c**). While model S02906 showed increase in epithelial features retaining substantial mesenchymal activity, model S02881_S2 adopted a particularly pronounced epithelial state at progression (cluster 3). Furthermore, E/M states as well as Hallmark EMT programs were evident in these progression-associated clusters (**Suppl. Fig. 9c**). These epithelial features were markedly diminished during subsequent tumor progression, suggesting that epithelial programs may facilitate the initial phase of tumor regrowth following a prolonged periods of tumor regression (**Fig. 6c, Suppl. Fig. 9c**).

In contrast, the chemotherapy-resistant model S02714 exhibited high concurrent epithelial and mesenchymal programs at baseline, suggestive of a more hybrid E/M-like state^57,58^. Following chemotherapy exposure, S02714 revealed expansion of a predominant tumor cell subpopulation (cluster 1) which was accompanied by a marked and comparable reduction in both epithelial and mesenchymal features following chemotherapy (**Fig. 6c, Suppl Fig. 9c**). Similar to the chemotherapy-sensitive models, resumed tumor progression at day 17 coincided with a transient upregulation of epithelial features and a modest increase in mesenchymal programs. Thus, despite the absence of measurable tumor shrinkage, chemotherapy treatment accomplished tumor growth control in the resistant model, and the transition to active tumor expansion coincided with the transient increase in epithelial features, similar to the chemotherapy-sensitive models. The maintenance of an overall balanced hybrid E/M state throughout treatment may, in turn, prime the resistant tumor for rapid regrowth following chemotherapy.

As all three PDX models were characterized by predominant expression of the lineage-defining transcription factors *ASCL1* and/or *NEUROD1* (**Fig. 1e, Supplementary Table 20**), we further investigated whether their expression changed throughout chemotherapy treatment. Across most tumor samples, we observed no major shifts in the overall distribution of lineage transcription factor expression or NE markers as well as unaltered *DLL3* expression, indicating overall stable SCLC subtype identity despite platinum-based treatment and tumor recurrence (**Suppl. Fig. 9d–e).** However, in the chemotherapy-sensitive model S02881_S2 transient tumor cell states emerging under therapeutic pressure were characterized by altered expression of *ASCL1* and NE markers. Although *YAP1* did not define a transcriptional subtype in any of our PDX models, we identified that these transient cell populations in S02881_S2 (cluster 3) – along with EMT-related signatures – exhibited markedly increased *YAP1* expression and associated YAP/TAZ downstream programs (**Fig. 6d**, **Suppl. Fig. 9f–g**). Further analysis of these transient cell states in S02881, revealed associations with basal-/progenitor-like features, indicating a plastic, stress-adapted state that may fuel into resumption of tumor cell growth (**Suppl. Fig. 9g**). In this model, transient modulation of SCLC subtype-defining lineage transcription factors without altering the overall subtype identity of the tumors at recurrence, may reflect a necessity to adapt to treatment-associated stress.

Together, our data suggest that chemotherapy-sensitive tumors survive treatment through pronounced but transient cell-state remodeling, with epithelial or hybrid E/M-associated states emerging around the onset of tumor regrowth, whereas resistant tumors preferentially rely on pre-existing adaptive states that are rapidly selected under therapeutic pressure. Notably, these dynamic adaptations occurred without stable/manifested conversion of the underlying SCLC lineage subtype.

## Discussion

Longitudinal tumor material from patients with SCLC is largely inaccessible, which limits comprehensive studies of therapy resistance in this deadly disease. Here, we leveraged a large in-house cohort of SCLC PDX models to systematically integrate controlled *in vivo* chemotherapy perturbations with genomic and single-cell transcriptomic profiling, enabling us to delineate how clonal selection, therapy-induced mutagenesis, and adaptive transcriptional programs jointly shape both response and resistance to platinum-based chemotherapy – which to date remains the cornerstone of SCLC treatment despite the emergence of promising novel treatment regimens^10,11^. We demonstrate that SCLC PDX models not only mirror patient treatment responses^2,32,34^ but also faithfully capture key tumor evolutionary processes observed in patients. Building on our previous comprehensive analysis of genome evolution in patients with SCLC^13^, we found similar evolutionary patterns recapitulated *in vivo*. Resistant tumor models largely retained stable clonal architectures with limited diversification under treatment, whereas chemotherapy-sensitive tumors underwent extensive clonal remodeling with frequent branching trajectories, depletion of dominant clones and expansion of pre-existing subclones. While prolonged *in vivo* propagation of tumor cells could introduce genetic drift, we and others have demonstrated stable genomic patterns across passages in these SCLC PDX models^34,59^, supporting that evolutionary changes observed here primarily reflect therapeutic selection. This is further evidenced by the detection of platinum-associated mutational signatures, which were present not only in models derived from chemotherapy-refractory patients, but were readily induced after only two cycles of *in vivo* chemotherapy in treatment-naïve models; consistent with observations in patients and supporting chemotherapy as a major selective pressure driving these evolutionary processes^60–62^. Furthermore, under sustained therapeutic pressure, recurrent alterations emerged in canonical SCLC-associated genes, including *TP73*, *MYC* family genes, chromatin regulators, and *NOTCH* pathway components, similarly identified in patients and further corroborating their relevance in driving tumor progression^13^.

Consistent with our previous findings in patient tumors^13^, we observed in our *in vivo* models that the common ancestral clone frequently persisted and constituted a substantial fraction of the recurrent tumor, emphasizing its critical role in withstanding therapeutic pressure and driving relapse. By integrating genome evolution with single-cell transcriptomics, we extend the utility of PDX models beyond our previous study^13^, enabling high-resolution characterization of the molecular states that emerge under therapeutic pressure. Our findings demonstrate that genome evolution is a major driver of transcriptional heterogeneity. Similarly, recent studies have linked ecDNA-mediated *MYC* amplification to transcriptional plasticity and therapy resistance^25,63^, while EMT states have been associated with genomic instability and tumor evolution^54^. Consistent with these observations, our integrative analyses identified a close association between platinum-associated genomic scars and EMT, including hybrid E/M states^57,58^. Model-based perturbation analyses across all single-cell datasets further emphasized EMT and selective epithelial and/or mesenchymal states as key determinants of the resistant phenotype across models. Notably, EMT-associated plasticity has repeatedly been implicated in therapeutic adaptation and resistance across different tumor entities and treatment modalities, including resistance in SCLC to platinum-based chemotherapy^23,64,65^ and DLL3-directed T cell engagement^66^ , EGFR-targeted therapies in NSCLC^67^ and chemotherapy in pancreatic, breast, ovarian cancer or cutaneous cell carcinoma^68–71^. This recurrent association may reflect the capacity of epithelial-mesenchymal plasticity to enable tumor cells to adopt alternative cell states that promote survival under therapeutic stress supporting distinct context-dependent adaptive phenotypes. Our findings extend these observations to platinum-treated SCLC, highlighting epithelial-mesenchymal plasticity as a potential adaptive mechanism under chemotherapy pressure.

Although previous studies have implicated NE changes and lineage switching in SCLC treatment resistance – based on limited patient samples and *in vitro* studies of murine SCLC models^22,26,27^–, our single-cell analysis across 96 tumors samples from 33 *in vivo* models did not support NE trans-differentiation as an obligatory feature of recurrent SCLC following platinum-based treatment. Across all models, *ASCL1*, *NEUROD1*, and/or *POU2F3* expression patterns were predominant and retained. Thus, our findings suggest that epithelial-mesenchymal plasticity can contribute to therapeutic adaptation without requiring stable changes in NE lineage identity^64,72^. Although *YAP1* expression was low across all tumors, one model showed transient YAP/TAZ activation coinciding with epithelial/EMT-associated transcriptional features during tumor regrowth, suggesting a potential interplay between Hippo/YAP/TAZ signaling and epithelial plasticity that may contribute to recurrence^73,74^. Notably, longitudinal tumor analysis demonstrated stable *DLL3* expression under chemotherapy pressure, supporting the rationale for combining platinum-based treatment with DLL3-directed therapies such as tarlatamab^10,11^.

The mechanisms by which residual tumor cells initiate regrowth following successful platinum-based chemotherapy remain an important and unresolved clinical question. To this end, longitudinal *in vivo* profiling of tumors under chemotherapy pressure revealed transient stress-associated transcriptional adaptations, including cell-cycle arrest and E/M activation, particularly increased epithelial features, that coincided with tumor progression after initial growth control and durable regression. These populations expanded under chemotherapy but declined upon recurrence, consistent with reversible persister-like states rather than stable lineage transitions^29,75^. Whereas our findings implicate that resistance is ultimately driven by genetically selected clones, these transient cell states appear to facilitate short-term survival under therapeutic stress; our work further emphasizes EMT as potential therapeutic vulnerability, warranting investigation of whether its inhibition can prolong tumor control and delay regrowth.

Together, our findings position chemotherapy resistance as a multi-layered evolutionary process driven by selection from ancestral clones, therapy-induced mutagenesis, and adaptive transcriptional programs among which EMT emerges as a central integrative axis that coordinates stress-response pathways without necessitating lineage reprogramming. From a translational perspective, the data underscore the importance of early intervention, targeting shared ancestral vulnerabilities, and limiting mutagenic therapeutic pressure to improve the durability of clinical responses.

**Supplementary Fig. 1.**
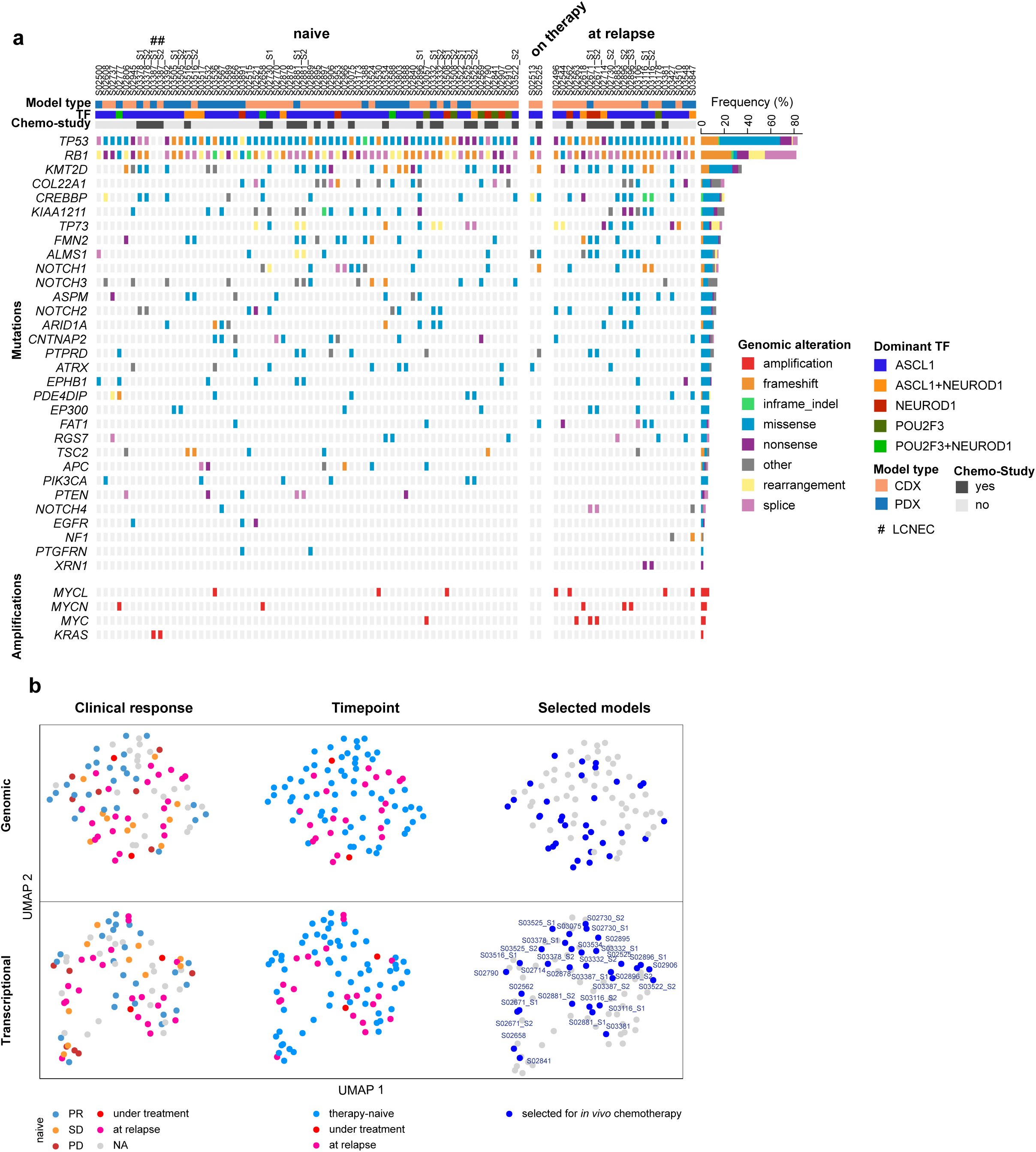
in support of Fig. 1 Molecular characterization of SCLC xenotransplant models and association with chemotherapy response. **a**, Key genomic alterations across 85 PDX models, focusing on selected SCLC-associated genes previously identified as significantly altered in genomic studies^13,15^. Genes are ranked from top to bottom according to alteration frequency. Models are grouped from left to right by ascending model ID and grouped by tumor sampling timepoint (treatment-naïve, during therapy and at relapse). **b,** UMAP embeddings of key genomic alterations (upper panel), as annotated in (a), and transcriptional profiles of 85 PDX models (referring to bulk RNA-Seq data). Models are colored according to clinical patient response (left panels) or tumor sampling timepoint (middle panels), and the 33 PDX models selected for *in vivo* chemotherapy treatment are highlighted (right panels).

**Supplementary Fig. 2.**
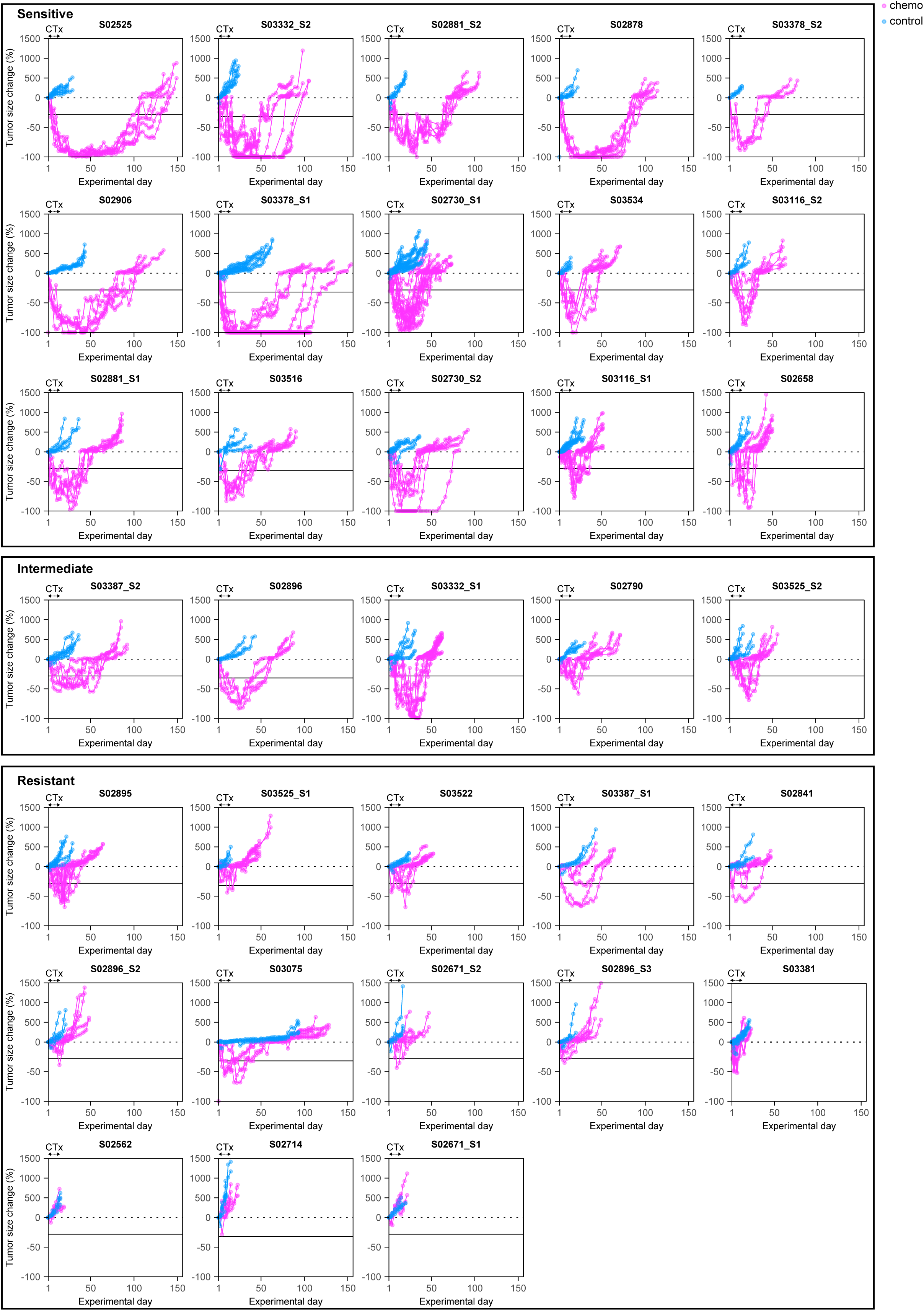
in support of Fig. 1 Relative tumor growth curves for individual xenotransplant models. Relative tumor growth curves (fold change from baseline) of 33 PDX models under chemotherapy (magenta) and control (blue) conditions. Each line corresponds to an individual tumor. The duration of chemotherapy treatment (CTx), corresponding to two cycles administered over 10 days, is indicated for each model. Models are stratified according to *in vivo* response groups, as defined by a composite score including best response and duration of response (**Methods, Supplementary Table 5**).

**Supplementary Fig. 3.**
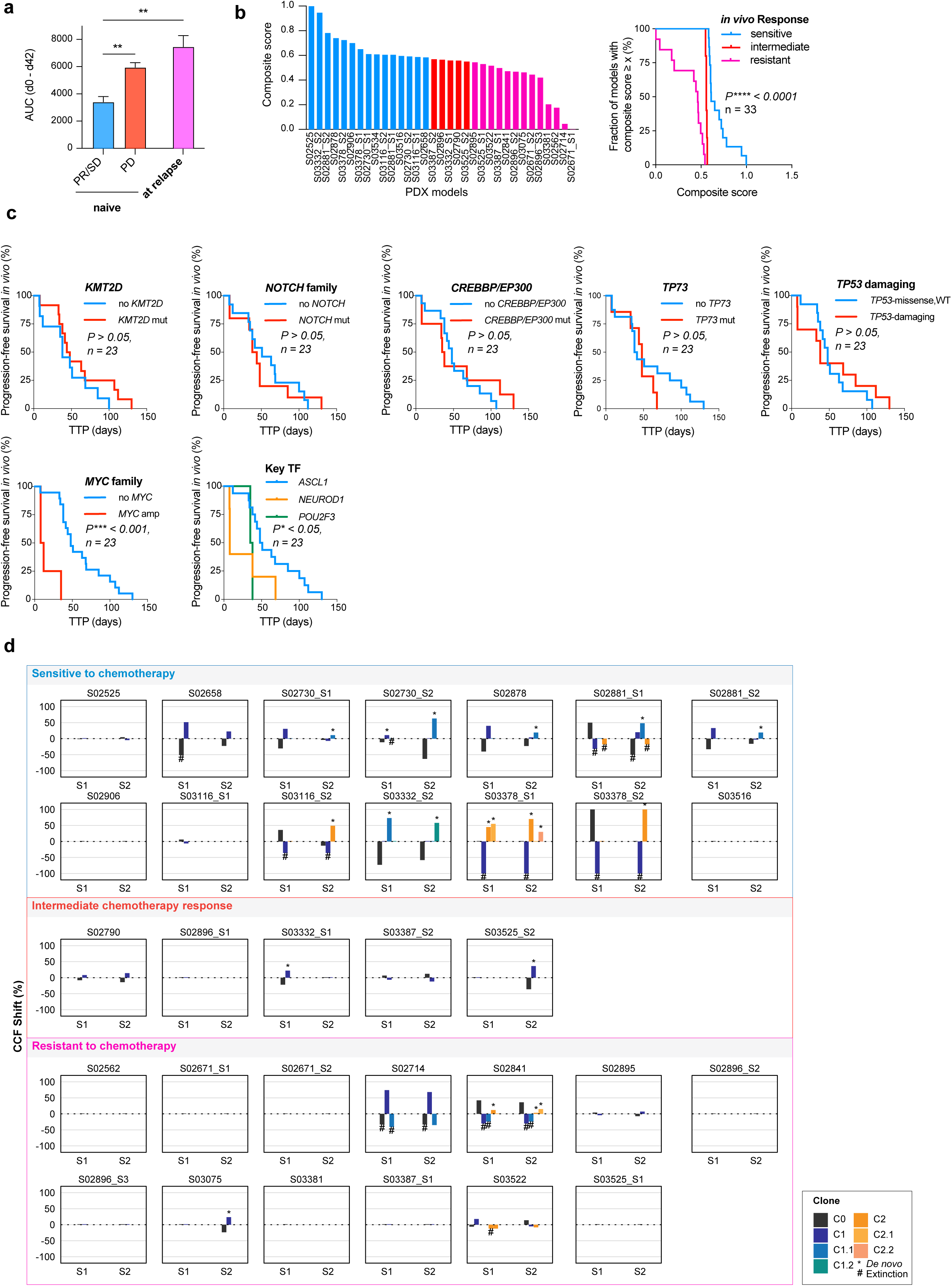
in support of Fig. 1 and 2 Chemotherapy response, molecular features and clonal dynamics. **a**, Box plot showing the area under the curve (AUC, days 0–42) of tumor growth in chemotherapy-treated PDX models (data shown in Fig. 1d), stratified by clinical patient response. Bars represent mean ± SEM. Unpaired two-tailed Welch’s t-test, \*\**P* < 0.01. **b,** Composite score integrating normalized duration of response and best response across 33 chemotherapy-treated PDX models (left). Kaplan-Meier curve showing the fraction of PDX models with composite scores exceeding the indicated threshold, stratified by *in vivo* response classification (right). Log-rank (Mantel–Cox) test, \*\*\*\**P* < 0.0001. **c,** Kaplan-Meier curves showing *in vivo* progression-free survival (TTP, days), stratified by *KMT2D*, *NOTCH* family, *CREBBP/EP300*, *TP73*, *TP53*-damaging and *MYC* family alterations or key transcription factor subtype. One PDX model per patient was included (n = 23; first model, prioritizing PDX over CDX). Log-rank (Mantel–Cox) test, \*\*\**P* < 0.001, \**P* < 0.05. **d,** Cancer cell fraction (CCF) shifts of individual SNV-defined clones between control and chemotherapy-treated tumors for each PDX model, stratified by chemotherapy response as sensitive, intermediate, or resistant. Bars indicate absolute changes in CCF of chemotherapy-treated samples relative to control. *De novo* clones are marked with an asterisk (*) and extinction of clones are marked with a hash (#). Clone identity is indicated by color.

**Supplementary Fig. 4.**
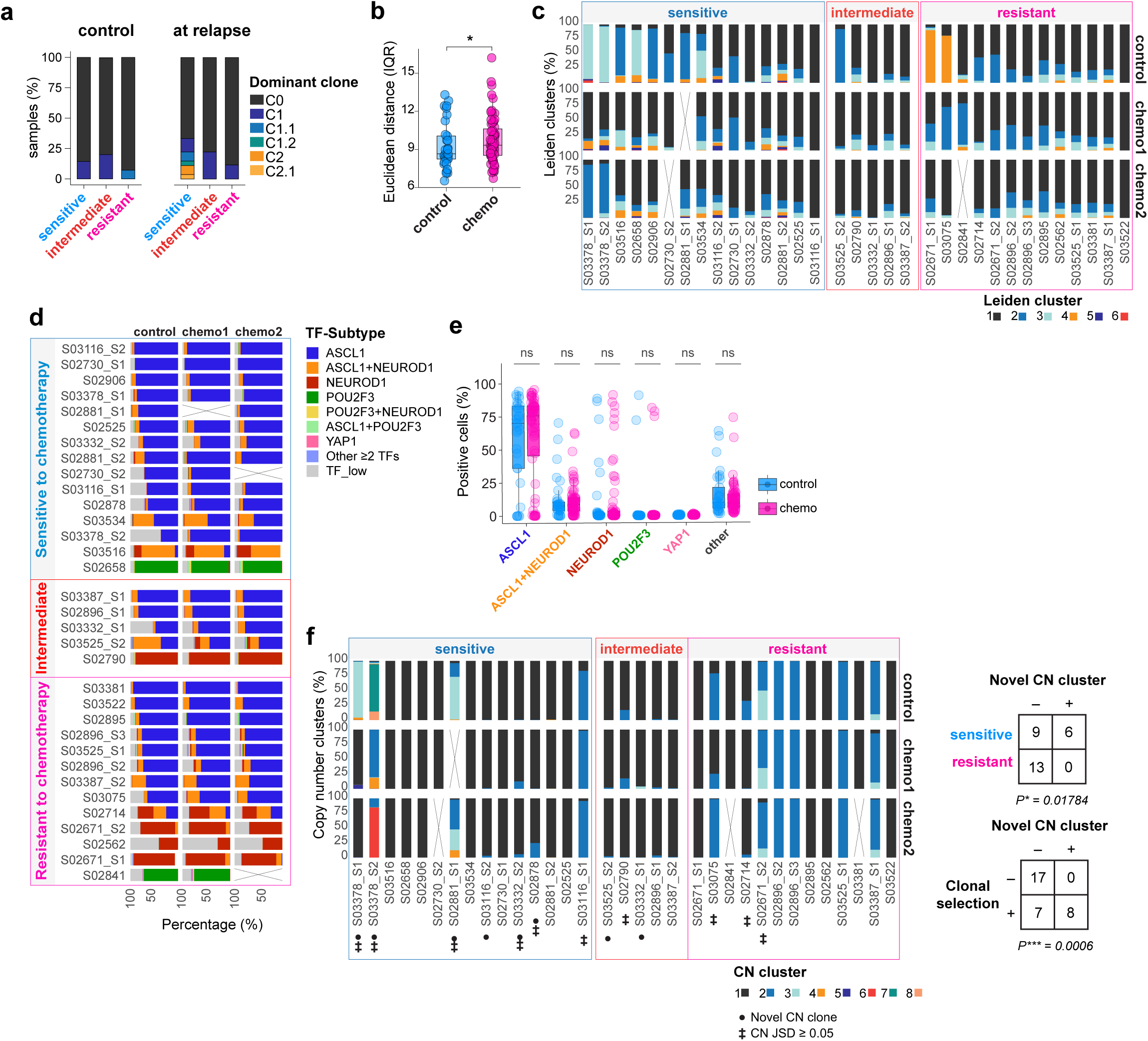
in support of Fig. 2 Genomic and transcriptional heterogeneity and evolution under chemotherapy. **a**, Distribution of the most dominant SNV-based tumor clones in control and chemotherapy-treated PDX samples at relapse, stratified by *in vivo* response group. **b,** Intratumoral transcriptional heterogeneity assessed from scRNA-seq data based on the interquartile range (IQR) of Euclidean distances between cellular transcriptional profiles within each sample, comparing control (n=33) and chemotherapy-treated tumors (n=63) across 33 PDX models. Individual samples are shown; biological replicates were averaged per model for two-sided paired Wilcoxon signed-rank test, *P*\*<0.05. **c,** Relative abundance of Leiden-defined transcriptional clusters identified from scRNA-seq data in control and chemotherapy-treated tumors across *in vivo* chemotherapy response groups. Crossed-out cases indicate missing biological replicates**. d,** Percentage of cells assigned to TF expression-defined subtypes based on gene-specific expression cutoffs for *ASCL1*, *NEUROD1*, *POU2F3* and *YAP1* in control (S1) and chemotherapy-treated tumors (chemo 1 and 2; S2 and S3) across 33 PDX models. “Other ≥2 TFs” includes all remaining combinations. Crossed-out cases indicate missing biological replicates. **e,** Percentage of cells assigned to TF expression-defined subtypes across control (n = 33) and chemotherapy-treated samples (n = 63). “Other” includes all remaining TF combinations (“Other ≥2 TFs” and “TF-low” cells). Individual samples are shown; biological replicates were averaged per model for two-sided paired Wilcoxon signed-rank test, ns, *P* > 0.05. **f,** Copy-number clusters inferred from scRNA-seq data describing CN-based composition of clones in control (S1) and chemotherapy-treated (chemo 1 and 2; S2 and S3, respectively) tumor samples across 33 PDX models grouped according to the in vivo chemotherapy response group. Crossed-out cases indicate missing biological replicates. Novel CN-based tumor clones emerging in chemotherapy-treated samples are marked with a solid circle. Models with a shift in CN-based tumor clones revealing along transcriptional inter-sample heterogeneity with JSD ≥ 0.05 are marked with ‡. Contingency tables show the association of novel emerging CN-based tumor clones with the respective in vivo chemotherapy response groups, and the association with SNV-based clonal selection procession. Fisher’s exact tests; *P\** < 0.05, *P*\*\*\*<0.001

**Supplementary Fig. 5.**
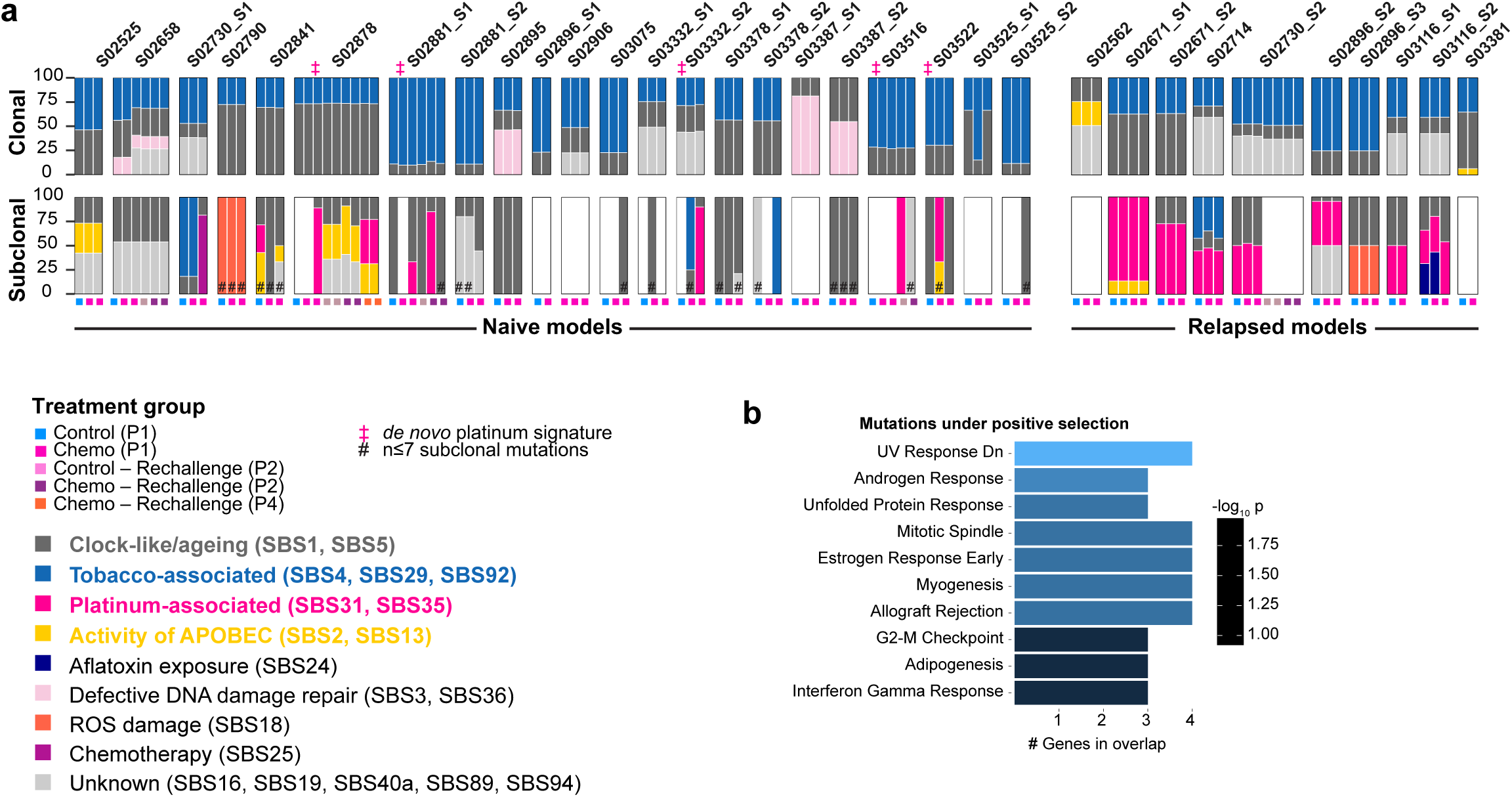
in support of Fig. 3 Mutational processes assigned to tumor clones across PDX tumors following *in vivo* chemotherapy treatment. **a**, Proportional contribution of mutational signatures to clonal and subclonal mutations across control- and chemotherapy-treated tumor samples for each model, grouped by treatment-naïve and relapsed PDX status. Treatment groups and mutational processes are color-coded. Samples with *de novo* acquired platinum-associated signatures and those containing ≤7 subclonal mutations are marked as specified. Blank spaces refer to tumor samples with no or very little sub-clonal mutations for which mutational signature assignment was not possile. **b,** Over-representation analysis of gene sets among mutated genes positively selected (with ΔCCF ≥ +30%) during *in vivo* chemotherapy treatment. Bars indicate the number of overlapping genes per gene set and are colored according to enrichment significance. Only genes expressed at FPKM ≥ 1 in at least 10% of our 85 PDX models were included.

**Supplementary Fig. 6.**
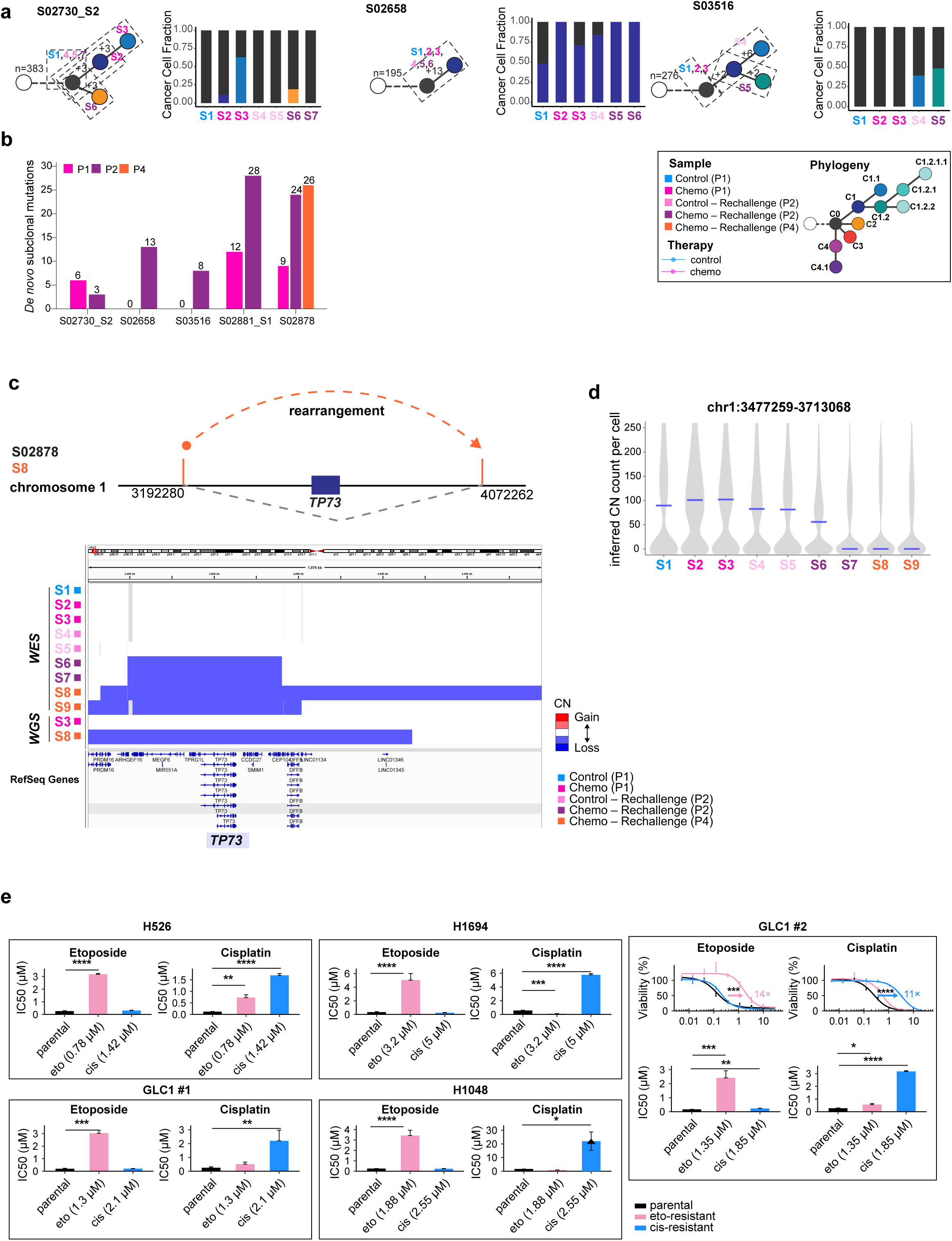
in support of Fig. 3 Genomic evolution during prolonged *in vivo* and *in vitro* chemotherapy exposure. **a**, SNV-based phylogenetic trees and corresponding CCF plots for tumor samples from PDX models retaining chemotherapy sensitivity during prolonged *in vivo* cisplatin-etoposide treatment (S02730_S2, S02658 and S03516). **b,** Number of *de novo* subclonal mutations detected across successive passages of rechallenged PDX models. **c,** Genomic rearrangement affecting *TP73* in model S02878. Top, schematic representation of the genomic rearrangement on chromosome 1 identified by WGS in sample S8, with breakpoints indicated in orange leading to loss of the region encompassing *TP73* (grey dashed line). Bottom, WES- and WGS-derived copy-number profiles across samples S1–S9, showing genomic loss emerging after P 2. **d,** Violin plots of scRNA-seq-inferred segment counts reflecting copy-number states at the *TP73* locus assigned for individual cells across longitudinal samples S1–S9 from S02878. Blue lines indicate median segment counts per sample. **e,** Acquired chemotherapy resistance following *in vitro* cisplatin or etoposide treatment of SCLC cell lines. IC_50_ values derived from CTG viability dose-response curves for cisplatin (cis) and etoposide (eto) are shown for parental and resistant derivatives of H526, H1694, GLC1 and H1048. Dose-response curves for GLC1 #2 are shown separately. Bars represent mean ± SEM. IC_50_ values exceeding the tested concentration range were capped at 30 µM for averaging and statistical testing, as indicated by triangles. Statistical significance was assessed using two-sided Welch’s t-tests on log_10_-transformed IC_50_ values. ns, P > 0.05; *\*P* ≤ 0.05; *\*\*P* ≤ 0.01; *\*\*\*P* ≤ 0.001; *\*\*\*\*P* ≤ 0.0001.

**Supplementary Fig. 7.**
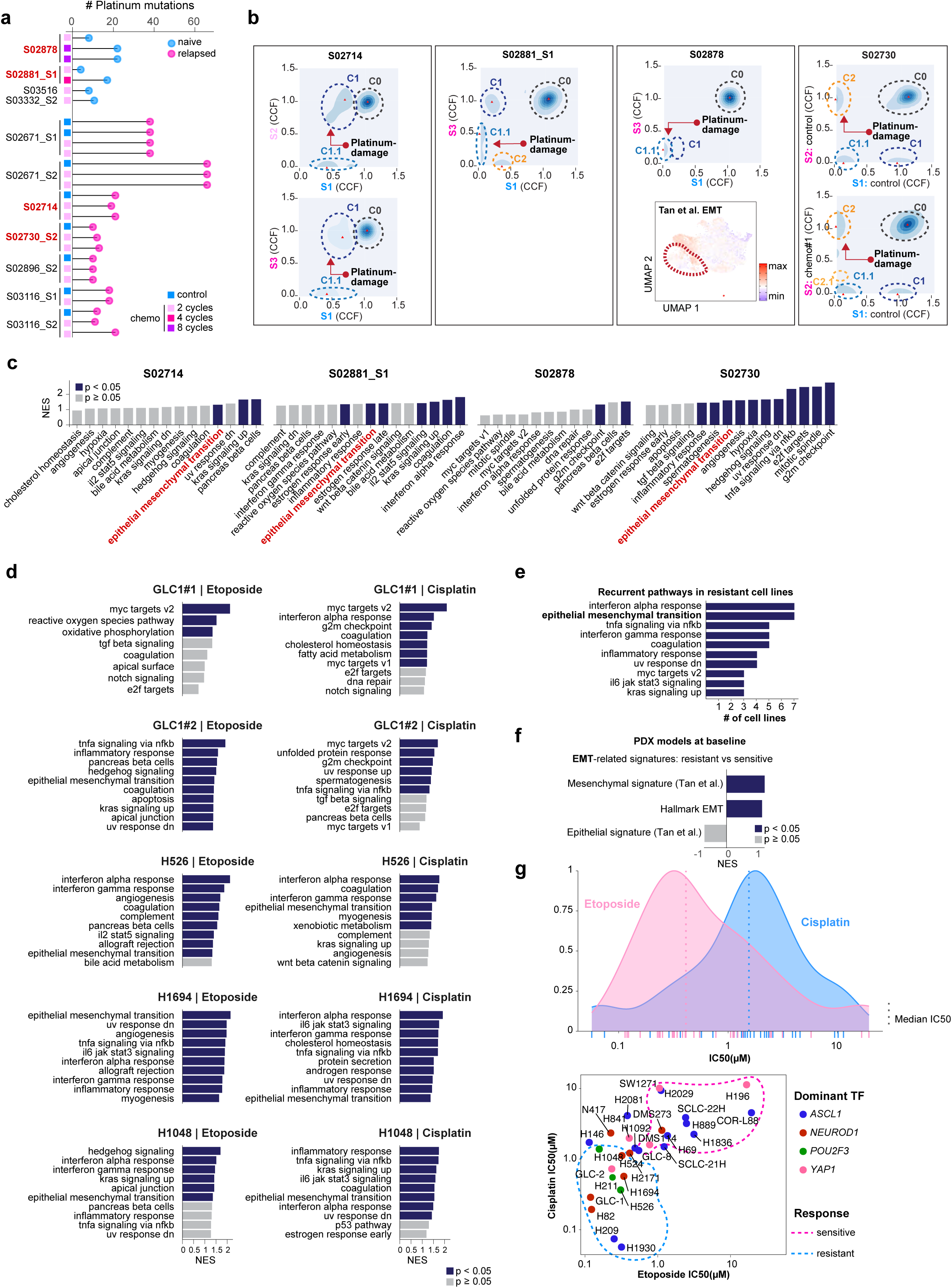
in support of Fig. 4 Characterization of platinum-associated clonal evolution and transcriptional programs linked to chemotherapy resistance. **a**, Number of platinum-associated mutations detected in individual tumor samples across n=11 PDX models following *in vivo* chemotherapy treatment. Treatment status of the corresponding patient (naïve or relapsed) and number of *in vivo* cisplatin-etoposide cycles are indicated. PDX models in which platinum-damaged clones could be tracked using scRNA-seq data (Fig. 4a) are highlighted in red. **b,** Representative CCF density plots of individual SNVs illustrating the corresponding clonal compositions across four PDX models (S02714, S02881_S1, S02878 and S02730). Clones harboring platinum-associated SBS signatures are indicated by red arrows. For S02878, EMT signature activity (Tan et al.^55^) is additionally projected onto the corresponding single-cell UMAP embedding. **c,** Gene set enrichment analysis (GSEA) of transcriptional programs associated with platinum-damaged cell populations identified through clone tracking shown in Fig. 4a. Bars indicate normalized enrichment scores (NES); significant pathways are highlighted in blue (P < 0.05). **d,** GSEA comparing baseline transcriptomic profiles of chemotherapy-resistant derivatives and their parental SCLC cell lines following prolonged *in vitro* exposure to etoposide or cisplatin. Top 10 Hallmark pathways are shown ranked by NES; significant pathways are highlighted in blue (P < 0.05). **e,** Recurrently enriched Hallmark pathways across chemotherapy-resistant cell line derivatives. Bars indicate the number of comparisons in which each pathway ranked among the top 10 positively enriched pathways (NES > 0, P < 0.05). Only pathways recurring in ≥3 comparisons are shown. **f**, GSEA of EMT-related signatures (Hallmark EMT and epithelial and mesenchymal signatures from Tan et al.^55^) in baseline PDX models comparing chemotherapy-resistant and chemotherapy-sensitive groups. Bars indicate NES; significant signatures are highlighted in blue (nominal *P* < 0.05). **g,** Distribution of cisplatin and etoposide IC_50_ values across SCLC cell lines (top), with median IC_50_ indicated as dashed lines. Scatter plot of cisplatin and etoposide IC_50_ values for individual cell lines (bottom), colored according to dominant transcription factor (TF) subtype. Dashed ellipses indicate chemotherapy-sensitive and -resistant cell line groups, defined using drug-specific median IC_50_ values.

**Supplementary Fig. 8.**
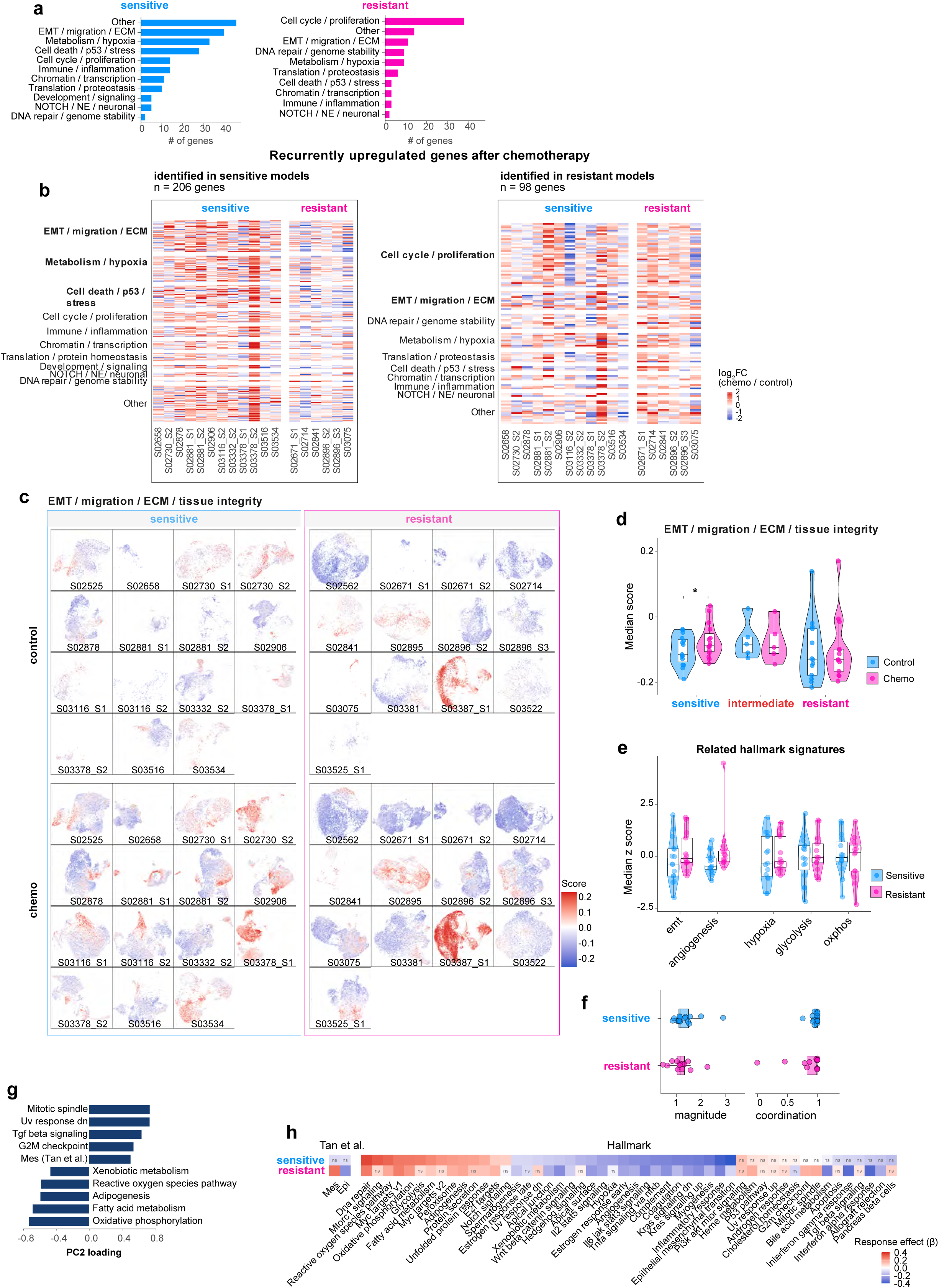
in support of Fig. 5 Response-specific transcriptional remodeling and cell states under chemotherapy. **a**, Functional annotation of recurrently upregulated genes identified by differential gene expression (DGE) analysis of condition-specific transcriptional clusters in chemotherapy-treated sensitive and resistant models. Genes were assigned to functional categories based on enrichment and annotation using Hallmark gene sets, Gene Ontology Biological Process (C5: GO:BP), and Reactome pathways. Bars indicate the number of genes assigned to each functional category, considering the primary functional category assigned to each gene. **b,** Heatmap of recurrently upregulated genes identified by DGE analyses of condition-specific clusters, grouped by functional category and displayed across sensitive and resistant models. For each sample, expression was aggregated across the selected condition-specific clusters, and log_₂_ fold changes between chemotherapy-treated and matched control tumors were calculated per model. Genes identified as recurrently upregulated in sensitive (left) or resistant (right) PDX models are displayed across both response groups and ordered by functional category. **c,** Single-cell projection of an EMT/migration/ECM/tissue integrity transcriptional score derived from recurrently upregulated genes identified by DGE of condition-specific clusters in chemotherapy-treated sensitive models. Scores are shown across control and chemotherapy-treated tumors from sensitive and resistant models. **d,** Median EMT/migration/ECM/tissue integrity scores in control and chemotherapy-treated tumors across chemotherapy-sensitive, intermediate and resistant response groups. Each point represents the median module score per PDX model and condition, with replicates pooled. Paired Wilcoxon signed-rank test; \*\**P* < 0.01 **e,** Z-scored median module scores of selected Hallmark gene sets associated with functional programs recurrently upregulated in chemotherapy-treated sensitive models, including EMT-associated programs (epithelial–mesenchymal transition and angiogenesis) and metabolic programs (hypoxia, glycolysis and oxidative phosphorylation), shown in baseline tumors from chemotherapy-sensitive and resistant PDX models. Two-sided Wilcoxon rank sum test, *P* > 0.05 **f,** CellOT-derived magnitude (L2 norm, global reference) and coordination (cosine similarity, within response group reference) of chemotherapy-induced transcriptional shifts in sensitive and resistant PDX models. Two-sided Wilcoxon rank sum test, *P* > 0.05. **g,** Top five positive and negative PC2 feature loadings from PCA of pathway activity scores, indicating the transcriptional programs most strongly associated with each direction of PC2. **h,** Heatmap showing response group-level effects (β) of CellOT-inferred chemotherapy-associated pathway shifts, estimated using linear mixed-effects models (LMMs) in sensitive and resistant PDX models. Positive and negative values indicate increased and decreased pathway activity following treatment, respectively. Statistical significance was assessed using two-sided Wald tests with Benjamini– Hochberg correction across pathways within each response group. Nonsignificant results (adjusted *P* ≥ 0.01) are labeled “ns”.

**Supplementary Fig. 9.**
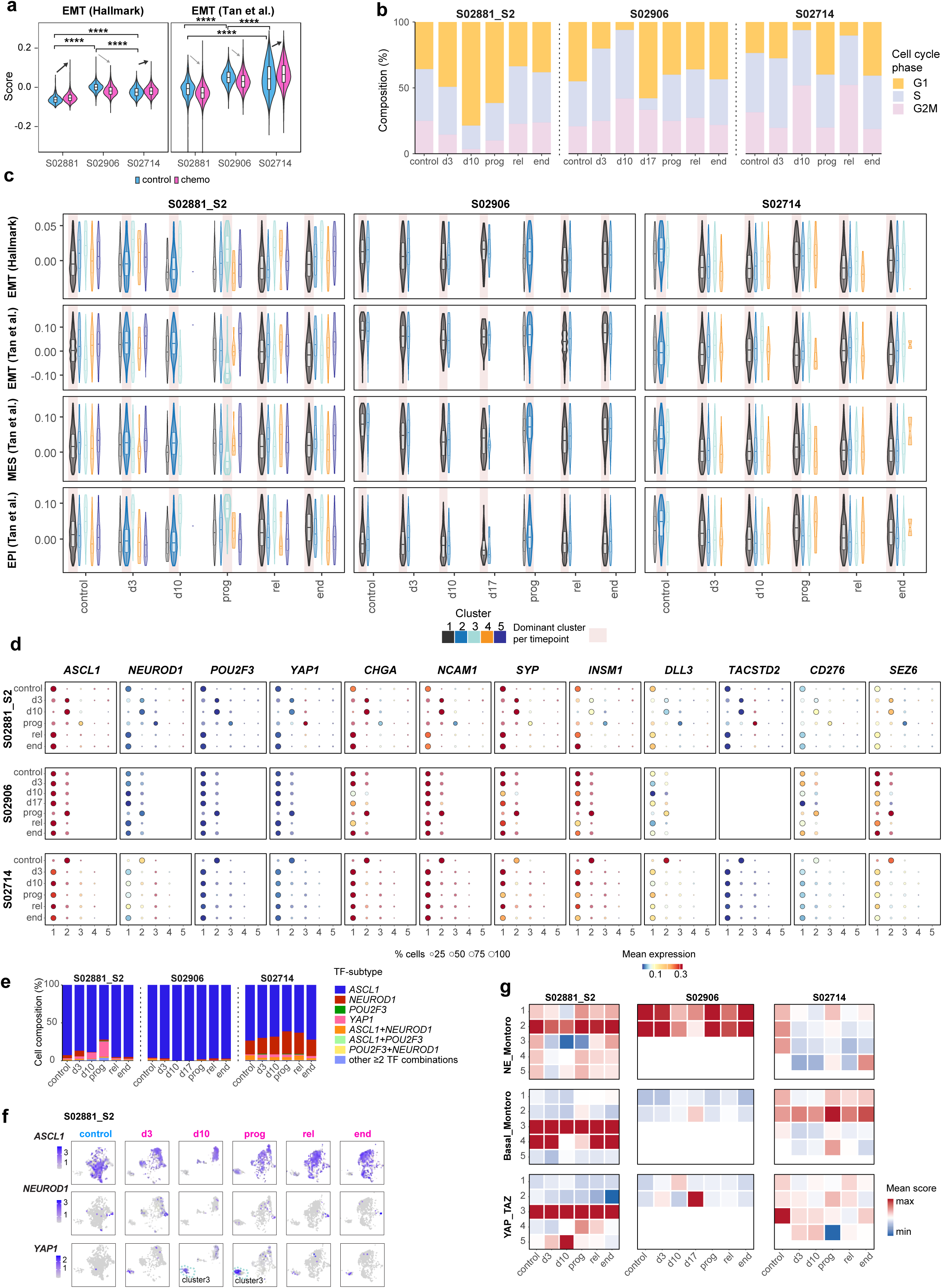
in support to Fig. 6 Longitudinal dynamics of EMT, cell-cycle and SCLC lineage programs under chemotherapy. **a**, EMT activity at the single-cell level in recurrent control and chemotherapy-treated tumors from S02881_S2, S02906 and S02714, assessed using scRNA-seq data and quantified based on the Hallmark EMT (left) and Tan et al. EMT (right) signatures. Statistical significance as indicated. One-sided Wilcoxon rank-sum test with BH-adjusted p-values; *\*\*\*\*P* < 0.0001. **b,** Cell-cycle composition across longitudinal sampling timepoints in chemotherapy-treated tumors from S02881_S2, S02906 and S02714, assessed by snRNA-seq data. Abbreviations: d, day; prog, progression; rel, relapse; end, endpoint. **c,** Distribution of Hallmark EMT, Tan et al. EMT^55^, mesenchymal (MES) and epithelial (EPI) signature scores^55^ across snRNA-seq-derived transcriptional clusters and longitudinal timepoints in the 3 PDX models. Colors denote individual clusters; the dominant cluster at each timepoint is highlighted. **d,** Expression of key SCLC lineage transcription factors and markers, including *ASCL1, NEUROD1, POU2F3, YAP1, CHGA, NCAM1, SYP, INSM1, DLL3, TACSTD2, CD276* and *SEZ6*, across snRNA-seq-derived transcriptional clusters and longitudinal samples. Dot size indicates the relative abundance of cells within each cluster per timepoint, and colors indicate mean expression. **e,** Composition of TF expression-defined SCLC cell states based on snRNA-seq data across longitudinal timepoints in S02881_S2, S02906, and S02714. TF subtypes were assigned based on gene-specific expression cutoffs for *ASCL1*, *NEUROD1*, *POU2F3*, and *YAP1*, determined separately from the snRNA-seq data. TF-low nuclei were excluded for better the visualization, proportions are shown for the remaining nuclei. **f,** UMAP projections of *ASCL1, NEUROD1* and *YAP1* expression across longitudinal timepoints in S02881_S2. **g,** Cluster-level activity of neuroendocrine^76^, basal^76^ and YAP/TAZ-associated transcriptional programs^77^ in snRNA-seq data across longitudinal sampling timepoints and clusters in S02881_S2, S02906 and S02714. Heatmap values indicate mean signature scores.

### Supplementary Tables

Supplementary tables will be published after peer-reviewed publication, and will be available upon request.

## Materials and Methods

### Animal experiments

#### Human tumor specimens

Tumor material was acquired at the University Hospital of Cologne and other collaborating hospitals as fine needle biopsies, lung resections, pleural effusions and circulating tumor cells (CTCs) following informed consent. Matching normal material was obtained as EDTA-anticoagulated blood samples. The study on patient material was approved by the institutional review board (IRB, reference: 19-1164) of the University of Cologne.

We analyzed a cohort of 85 patient-derived xenotransplantation (PDX) models across 71 patients established predominantly from treatment-naïve samples (n=62, 73%), with additional models derived from patients undergoing treatment (n=2, 2%) or at tumor recurrence following first-line therapy (n=21, 25%; **Supplementary Table 1**). Patients had a median age of 68 years at diagnosis (IQR 62–74), 73% had a reported history of smoking, and the majority presented with advanced-stage disease (UICC stage III–IV). Histological assessment of both the corresponding patient tumors and xenotransplant models was performed by expert pathologists (AQ, RB) at the University of Cologne (**Supplementary Table 19**). All models were classified as SCLC, with the exception of two models established from one patient (S03387) exhibiting large-cell neuroendocrine carcinoma (LCNEC) histology. Genomic profiling further supported the histological classification, demonstrating characteristic SCLC-associated genomic alterations, including *TP53* and *RB1* alterations and other recurrent SCLC-associated events (**Suppl. Fig. 1a, Supplementary Table 3**). In contrast, the two LCNEC models lacked *RB1* alterations and harbored *KRAS* amplifications.

A subset of 33 models was selected for functional *in vivo* treatments with cisplatin and etoposide (**Fig. 1a**). Histological assessment of these models confirmed SCLC morphology in n=31 models, and LCNEC histology in two models derived from the patient S03387. Of the 33 models, n=22 were established from treatment-naïve patients and n=10 from patients at relapse following first-line therapy. One additional model (S02525) was established from CTCs collected during the treatment period, specifically before initiation of first-line chemotherapy but shortly after the patient had received brain radiotherapy. As tumor cells were acquired before systemic chemotherapy exposure and no evidence of treatment-associated genomic alterations was detected, this model was classified as treatment-naïve for the purposes of this study. For 27 models derived from n=20 patients with SCLC (n=17 PDX models in the treatment-naïve setting), clinical response data following first-line platinum-etoposide-based chemotherapy were available and incorporated into the analysis. Clinical responses to platinum-based chemotherapy were classified as partial response (PR), stable disease (SD), or progressive disease (PD), with an additional category comprising models established from tumor cells at relapse (n=10). For the n=6 remaining models, including two models derived from the patient with LCNEC histology, clinical response data was not available. Clinical, histopathological and molecular characteristics for all PDX models are summarized in **Supplementary Table 1 and 19**.

#### Generation of patient-derived xenotransplant models

All animal experiments were conducted under specific-pathogen-free conditions in individually ventilated cage systems with a 12-h light/12-h dark cycle and controlled temperature and humidity. All experiments were approved and performed in accordance with the regulations of the local animal welfare authorities (Landesamt für Verbraucherschutz und Ernährung Nordrhein-Westfalen, AZ: 84–02.04.2018.A002 and 81–02.04.2022.A394).

To expand limited tumor material for molecular and functional studies, patient-derived xenograft (PDX) or circulating tumor cell-derived xenograft (CDX) models were established by subcutaneous engraftment into the flanks of 7–14-week-old female NOD.Cg-*Prkdc^scid^ Il2rg^tm1Wjl^*/SzJ mice (NSG mice, The Jackson Laboratory). Tumor tissues or cells isolated from pleural effusions were washed in PBS; pleural effusion samples were additionally depleted of erythrocytes using Red Blood Cell Lysis Solution (Miltenyi Biotec). Samples were mixed with Matrigel Matrix (Corning) and subcutaneously transplanted. For CDX models, CTCs were isolated as previously described^32^. Briefly, CTCs were enriched from at least 10 ml EDTA-anticoagulated blood using RosetteSep CTC Enrichment Cocktail (STEMCELL Technologies) followed by Ficoll-Paque^TM^ PLUS (Cytiva) density-gradient centrifugation. Isolated CTCs were washed, resuspended in HBSS:HITES (9:1), mixed 1:1 with Matrigel, and transplanted subcutaneously. Mice were monitored for tumor growth at least 1x weekly, and sacrificed when tumor volume reached a maximum of 1500 mm^3^, calculated as tumor length × width^2^/2, or earlier if signs of poor health were observed.

Harvested tumors were dissected into fragments for cryopreservation in FBS containing 10% DMSO, fresh-frozen (FF) archiving for downstream genomic and transcriptomic analyses, or for formalin fixation and paraffin embedding (FFPE) for histological studies. Histological analyses were performed by experienced pathologists at the Department of Pathology, University Hospital Cologne (AQ, RB), confirming tumor histology and a tumor content of at least 80% (**Supplementary Table 19**). Additionally, tumor tissue was enzymatically dissociated into single-cell suspensions using the gentleMACS system (Miltenyi Biotec) according to the manufacturer’s instructions. For *in vivo* propagation, tumor cells were engrafted into 1–2 NSG mice. All xenograft models underwent at least one passage before use in subsequent *in vivo* experiments.

#### Chemotherapy treatment *in vivo*

For controlled *in vivo* treatment studies, one million tumor cells derived from PDX or CDX models were engrafted subcutaneously into both flanks of 7–14-week-old NSG mice. Once tumors reached a volume of 100–200 mm³, mice were randomized to chemotherapy (3 mice, 5–6 tumors) or vehicle control (2–3 mice, 4-6 tumors). The chemotherapy group received intraperitoneal injections of cisplatin (5 mg/kg) on days 1 and 8, and etoposide (10 mg/kg) on days 1–3 and 8–10. Drugs were prepared by the Central Cytostatic Preparation Unit, Pharmacy, University Hospital Cologne, and diluted in PBS before administration. Control mice received PBS at the corresponding time points. Tumor volume was measured by caliper 3 times weekly. Mice were euthanized and tumors harvested for downstream analyses when tumor volume reached the predefined endpoint of 1,500 mm³ or earlier if signs of poor health necessitated termination.

Relative tumor growth was calculated as the percentage change in tumor volume relative to baseline, with tumor volume at treatment initiation (day 0) serving as the reference (100%, **Suppl. Fig. 2, Supplementary Table 4**). To visualize response dynamics according to the corresponding patient’s clinical response (PR/SD [n = 14], PD [n = 3], relapse [n = 10], NA [n=6], **Fig. 1d**), locally weighted scatterplot smoothing (LOWESS, 10-point window) was applied to all data points (n = 616; x-range 0–153 days) using GraphPad Prism. Tumor growth was further quantified by calculating the area under the tumor growth curves (AUC). Models were additionally assessed for tumor shrinkage, with a reduction in tumor volume of ≥30% from baseline defining a response. Chemotherapy response was further quantified as “best response” (BR), defined as the maximum percentage change in tumor volume from baseline after day 14 (following completion of two chemotherapy treatment), and as normalized “duration of response” referring to the normalized time to progression (TTP), defined as the number of days required for tumor volume to double relative to baseline and normalized to the mean tumor-doubling time of the corresponding vehicle group^34^. For comparative classification of the *in vivo* chemotherapy response across all models, BR and TTP were normalized to a 0–1 scale using min–max scaling. Since lower BR values indicate better responses, BR was inverted after min-max scaling. A composite score was calculated as the equally weighted mean of both normalized BR and TTP.

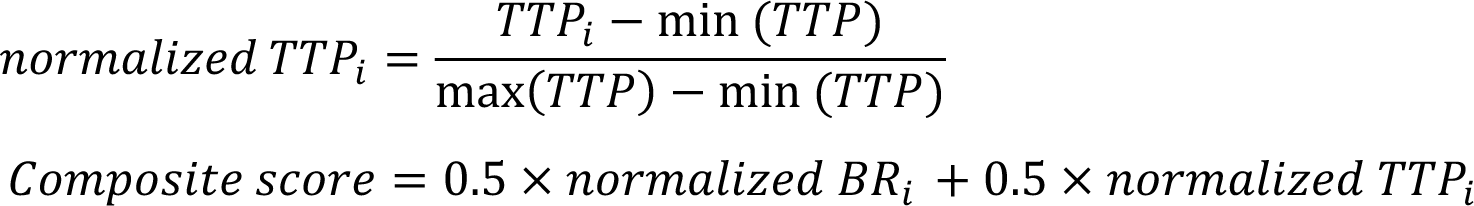

Thus, models were classified as chemotherapy-“sensitive” (good responders) for composite scores greater than 0.59, “intermediate” (moderate responders) for scores of 0.55-0.59, and chemotherapy-“resistant” (poor responders) for scores below 0.55 (**Fig. 2b**, **Suppl. Fig. 2, Supplementary Table 5**).

*In vivo* rechallenges of tumors with chemotherapy were performed for 5 chemotherapy-sensitive models (**Fig. 3b**). Tumors from chemotherapy-treated mice were processed and one million cells were subcutaneously reimplanted into both flanks of four 7–14-week-old female NSG mice per model. Once tumors reached ∼100–200 mm³, mice were subsequently randomized to additional chemotherapy or vehicle control treatment, using the same regimen described above. The serial transplantation and treatment strategy enabled a cumulative *in vivo* exposure of 4–8 cycles of chemotherapy across passages P2–P4, with longitudinal monitoring of tumor growth for up to 8 months. Tumor growth was monitored throughout treatment period, and tumors were collected for downstream analyses upon reaching the predefined termination criteria. Details of the tumor samples collected during these *in vivo* rechallenge experiments and the corresponding PDX models are provided in **Supplementary Table 2**.

To characterize treatment-induced tumor responses longitudinally – rather than only at the endpoint – an *in vivo* chemotherapy time-course study was performed in three PDX models (1 chemotherapy-resistant and 2 chemotherapy-sensitive models, **Fig. 6a**). For each model, 1 million tumor cells were engrafted subcutaneously into both flanks of 8–10 female NSG mice aged 7–14 weeks. Once tumors reached a size of 100–200 mm³, two mice were assigned to the vehicle control group, and the remaining mice received the cisplatin–etoposide regimen described above. Chemotherapy-treated mice were euthanized at predefined timepoints corresponding to day 3 (after completion of one treatment cycle) and day 10 (completion of two cycles), and at day 17 if tumors remained measurable. Subsequent sampling was determined by tumor growth kinetics, which were monitored three times weekly. One sample was collected at progression, defined as the first consistent increase in tumor volume across at least two consecutive measurements following a lower tumor volume plateau or complete regression. Further samples were collected when tumor size returned to the baseline volume at treatment initiation and when tumors reached the maximum permitted volume of 1,500 mm^3^. For vehicle-treated mice, tumors were harvested when they reached the endpoint of 1,500 mm^3^. All tumors were snap-frozen in liquid nitrogen immediately after collection and stored for subsequent nuclei extraction.

#### Nucleic Acid Extractions

Genomic DNA and RNA were extracted from 20 μm sections of fresh-frozen (FF) tumor tissue or from tumor cell pellets; DNA was additionally isolated from EDTA-anticoagulated blood. FF tissue sections were prepared using a cryostat (Leica) maintained at −20 °C. DNA was isolated using the Gentra® Puregene® DNA Purification kits (Qiagen) according to the manufacturer’s instructions. All DNA isolates were hydrated in TE buffer (Invitrogen) and quantified using PicoGreen (Thermo Fisher Scientific) before downstream applications. RNA was extracted using the RNeasy Mini Kit (Qiagen) according to the manufacturer’s instructions, eluted in 30–50 μl RNase-free water, quantified using a Nanodrop spectrophotometer (Thermo Fisher Scientific), and stored at −80 °C until further use. DNA and RNA integrity were assessed using the Agilent TapeStation system with Genomic DNA ScreenTape and RNA ScreenTape, respectively (Agilent Technologies). FF-derived DNA samples with high molecular weight (>10 kb) were used for whole-exome and whole-genome sequencing (WES and WGS), whereas samples showing substantial degradation were excluded. Samples with an RNA integrity number (RIN) above 7 were used for RNA sequencing (RNA-seq).

#### STR validation of tumor tissue and cell lines

Short tandem repeat (STR) profiling was performed to confirm the patient origin of xenograft-derived tumor tissues and matched normal blood samples. STR profiling was conducted at either the Institute of Legal Medicine, University Hospital of Cologne, or the Cologne Center for Genomics (CCG). Established cell lines were routinely authenticated by STR profiling, including prior to sequencing.

#### DNA sequencing and genomic analysis

DNA sequencing was performed using WES and WGS on Illumina NovaSeq or HiSeq platforms. WES libraries were prepared using the Twist Human Core Exome Kit (Twist Bioscience), and sequenced with 2×100-bp paired-end reads. WGS libraries were generated using the TruSeq DNA Nano Kit (Illumina) and sequenced with 2×150-bp paired-end reads at 30–60× coverage. All procedures followed the manufacturers’ instructions.

Raw sequencing reads were processed using our in-house pipeline *peiflyne* (http://www.uni-koeln.de/med-fak/peiflyne/peiflyne.tgz), as previously described ^13–15,78^. Reads from tumors collected from PDX/CDX models were aligned to a combined human–mouse reference genome (GRCh37/hg19 and GRCm38/mm10) to distinguish human tumor reads from murine reads and to exclude murine reads from downstream analyses. Patient-matched normal samples were processed similarly to ensure consistency. Sequencing data from chemotherapy-treated human SCLC cell lines were aligned to GRCh37/hg19, with the corresponding untreated parental cell line serving as a normal reference (**Supplementary Table 2**). Somatic mutations were identified, and somatic copy number alterations were assessed using *Sclust cn* ^78,79^ to estimate tumor purity, ploidy and allele-specific copy number states based on read depth and allelic imbalance. CNV profiles were inspected using Integrative Genomics Viewer (IGV^80^ v2.8.0). Median genome ploidy was 3.1 (IQR: 1.9–3.29). Across WES data from CDX/PDX models, mean sequencing coverage was 186× (interquartile range: 151–215×), and median tumor purity was 99%, due to the exclusion of non-human infiltrating cells and the resulting enrichment of human tumor-derived cells. Thus, our data provided sufficient depth and tumor purity for tumor-specific variant detection and clonal fraction estimation (see below). Of the 33 PDX models included in the *in vivo* chemotherapy study, 32 were included in WES-based analyses. One model (S03534) was excluded due to contamination with another tumor, which could not be reliably filtered out from the WES data. Additional WGS was performed on selected tumor samples to characterize complex genomic alterations and structural rearrangements (**Supplementary Table 2**).

#### Bulk-RNA sequencing and expression analysis

Bulk RNA sequencing was performed using poly(A)-selected messenger RNA (mRNA) to generate cDNA libraries with the Illumina TruSeq mRNA Library Preparation Kit according to the manufacturer’s instructions. Libraries were sequenced on Illumina NovaSeq or HiSeq platforms with 2×100-bp paired-end reads, yielding approximately 50 million reads per sample, corresponding to a mean coverage of 30× across the annotated transcriptome.

Data were processed using the Transcriptome Reconstruction and Unified Pipeline^81^ (*TRUP*). For PDX/CDX models, reads were aligned using STAR^82^ to a combined human-mouse reference genome (GRCh37/hg19 and GRCm38/mm10). Human SCLC cell line reads were aligned to GRCh37/hg19. Expression levels were determined with Cufflinks version 2.2.1^83^ for paired-end reads uniquely mapped to the human reference genome, quantified as fragments per kilobase of exon per million mapped fragments (FPKM) and as gene-level read counts. RNA-Seq data for parental SCLC cell lines were obtained from previously published datsets generated by Genentech^16^, or from our work^21,84^. Expression levels in these datasets were quantified using *kallisto* v0.44.0^85^ and reported as transcripts per million (TPM).

RNA-seq data for 85 PDX/CDX models (**Fig. 1b, Suppl. Fig. 1b**) were examined using Uniform Manifold Approximation and Projection (UMAP) based on variance-stabilized (VST) counts of the 1,000 most variable genes (*uwot* v0.2.3). Differential expression analysis of PDX/CDX models and cell lines was performed with *DESeq2* (v1.42.1). Pathway enrichment was assessed using pre-ranked *fgsea* (v1.28.0) based on log₂ fold changes against the MSigDB Hallmark collection^53^.

#### Clonal architecture and phylogenetic reconstruction

Tumor clonality analyses and phylogenetic reconstructions were based on somatic single-nucleotide variants (SNVs) identified by WES of multiple patient-matched tumor samples, using *Sclust*^78^ and previously described methods^13,86^. In brief, for each mutation in each sample, an expected allelic fraction was determined, correcting for tumor purity, average ploidy and the local copy number state. Cancer cell fractions (CCFs) were then estimated by comparing expected and observed allelic fractions. To reduce sequencing noise, *peiflyne mfilter* was applied using sample-specific read-count thresholds (10–30 reads), excluding low-frequency variants attributable to sequencing noise while retaining well-supported subclonal mutations.To reconstruct clonal relationships and infer patient-level phylogenies, all available samples from each patient, including samples from multiple PDX/CDX models where applicable, were jointly re-processed and analyzed. Copy number profiles were harmonized using consensus segmentation with shared breakpoints across all patient-matched samples to determine allele-specific copy numbers. Somatic variants were re-queried across matched samples to recover low-frequency mutations. Quality filtering excluded variants affected by mapping artifacts (including human–mouse cross-mapping), unreliable copy-number states, loss of heterozygosity, segmentation gaps, or insufficient or excessive sequencing coverage. A binomial non-detection test with false discovery rate (FDR) correction was applied to distinguish private mutations from shared mutations that were not detected in all samples. Filtering criteria were harmonized across all patient-matched samples to ensure consistent mutation assignments, followed by model-specific clonal assignments and phylogenetic reconstruction. We thus assigned a common ancestral clone (C0), characterized by cluster of mutations with CCF of 100%, and descendant subclones (C1, C2, C3, as illustrated in **Fig. 2a**), referring to clusters of mutations that were either private to individual samples or shared between specific subsets of samples. Tumor clones were identified by pairwise two-dimensional CCF clustering using *Sclust cluster2d*, based on smoothed two-dimensional CCF distributions. Shared and private mutations were assigned to their respective clusters (with at least 3 mutations required to define a tumor clone). Clustering parameters (*lambda* and *nbins*) were adjusted to optimize cluster separation, and ambiguous mutation assignments were resolved or excluded. Clonal phylogenies were reconstructed under the infinite-sites assumption, the CCF sum rule, and the “once private, never shared” constraint^13,87^. When multiple solutions were possible, linear evolutionary trajectories were favored. For uniquely resolved phylogenies, cluster-level CCFs were normalized to a shared patient-level common ancestral clone (C0) and decomposed into clone-level CCFs.

To assess chemotherapy-induced clonal selection, comparisons were restricted to control and chemotherapy-treated samples within each model. Models were classified as exhibiting “*Selection”* if treatment resulted in clonal extinction (loss of subclones detected in control but not treated samples), *de novo* emergence of treatment-specific subclones, or expansion of pre-existing subclones (with ΔCCF ≥ 30%); models showing none of these changes in clonal dynamics were classified as *“No selection”* (**Fig. 2a, 2d, Suppl. Fig. 3i**). Single-cell copy-number profiles were additionally considered to assess genomic changes (**Fig. 2g, Suppl. Fig. 3h**). Genes mutated in expanding subclonal mutations were subjected to pathway enrichment using *EnrichR* v3.4, restricted to genes expressed in ≥10% of xenograft models (log₂(FPKM + 1) ≥ 1, **Suppl. Fig. 5b**).

#### Mutational signature analysis

Mutational signature analysis was performed using SigProfiler^43,88^. Somatic SNVs identified by WES were converted into 96-channel single-base substitution (SBS-96) matrices using SigProfilerMatrixGenerator v1.2.26. Contributions of known mutational signatures from the COSMIC v3.4 reference catalogue^89^ were estimated using non-negative least squares (NNLS) optimization. Analyses were performed separately for clonal and subclonal mutations passing quality filters for clonal cluster assignments (see above). For treatment-naïve models, clonal status was defined per model, whereas for models established after clinical relapse, it was defined across all available models from the same patient (PDX and CDX) to better resolve ancestral and therapy-associated subclonal mutations acquired during clinical treatment. To minimize overfitting and noise, signatures were initially fitted against the full COSMIC catalogue^89^, followed by selection of signature subsets based on recurrence and contribution (**Supplementary Table 13**). For clonal mutations, signatures detected in more than one patient and represented by at least 20 mutations per sample were retained. Of the 12 signatures meeting these criteria (SBS5, SBS4, SBS3, SBS40a, SBS92, SBS2, SBS13, SBS16, SBS29, SBS36, SBS89, SBS8), SBS8 was excluded because of its uncertain etiology, low prevalence and potential ambiguity in signature attribution. SBS1 was added because of its relationship to SBS5, resulting in a final set of 12 signatures. For subclonal mutations, signatures were retained if detected in at least one sample with more than 20 subclonal mutations and accounting for at least 30% of mutations in that sample. Of the 12 signatures meeting these criteria (SBS31, SBS5, SBS35, SBS7b, SBS13, SBS18, SBS19, SBS24, SBS25, SBS29, SBS94, SBS99), SBS7b was excluded because of its UV-associated etiology and similarity to SBS31. Four additional biologically related signatures were manually added based on their etiological similarity to the detected signatures (SBS1, SBS2, and SBS92) and to account for tobacco-associated signatures (SBS4) which were highly prevalent among clonal mutations. This resulted in a final set of 15 signatures. Signature fitting was then repeated separately for clonal and subclonal mutations using the respective signature subsets. Mutational signature contributions were assessed in control and chemotherapy-treated PDX samples (**Supplementary Fig. 5a**), with contributions within post-treatment samples specifically displayed in **Fig. 3a**. Mutational signatures assigned to samples with low subclonal mutation counts (≤7 mutations) are specifically flagged in the analyses due to increased uncertainty in the reliability of signature attribution (**Suppl. Fig. 5a**). Analyses were also performed in SCLC cell lines following *in vitro* exposure to either cisplatin or etoposide (**Fig. 3d**). The assignment of mutational signatures across all samples is summarized in **Supplementary Table 13**.

#### Single-cell and single-nuclei RNA sequencing (scRNA-seq, snRNA-seq)

Single-cell RNA sequencing (scRNA-seq) was performed on viable tumor tissues harvested from control and chemotherapy treated tumors at the experimental endpoint. For each model, one control tumor (S1) and two chemotherapy-treated tumors (chemo 1 and chemo 2, S2 and S3, respectively) were collected from separate mice, and processed within 1–2 h of harvest. Single cell suspensions with a viability of at least 75% were adjusted to 1,000 cells/μl in PBS containing 0.04% bovine serum albumin (BSA), and up to 10,000 cells were loaded onto the 10x Genomics Chromium platform. Samples with lower viability underwent dead-cell removal by Ficoll density-gradient centrifugation, with the procedure repeated until the required viability was achieved. The Chromium Next GEM Single Cell 3ʹ Reagent Kit v3.1 (10X Genomics) was used to generate partitioned single-cell cDNA and library preparation according to manufacturer’s instructions. Overall, scRNA-seq data were generated for 33 PDX models following chemotherapy treatment, comprising 33 control and 63 chemotherapy-treated tumor samples. Notably, processing of viable cells and subsequent single-cell library preparation were unsuccessful for biological replicates in three models, leaving only one chemotherapy-treated tumor sample available for analysis in each of these models (**Supplementary Table 2**).

For the longitudinal PDX/CDX experiment (**Fig. 6**), single-nucleus sequencing (snRNA-seq) was performed on flash-frozen tumor tissues (1–3 per timepoint) because viable tumor cell isolation was not feasible for samples collected under chemotherapy. Nuclei were isolated from snap-frozen tumor tissues stored at −80 °C. All procedures were performed on ice or at 4 °C to minimize RNA degradation. Frozen tissue was lysed in 2 ml ice-cold Nuclei Extraction Buffer (Miltenyi Biotec) supplemented with 0.2 U/μl murine RNase inhibitor (New England Biolabs) and mechanically dissociated using a gentleMACS dissociator according to manufacturer’s instructions. Nuclei were stained with DAPI (Miltenyi Biotec) and sorted by flow cytometry using a BD FACSAria IIIu or a BD FACSAria Fusion Cell Sorter (BD Biosciences, San Jose, CA, USA) with 70 μm nozzle at 70 psi pressure. Single nuclei were detected based on their DAPI fluorescence and debris, aggregates and non-intact nuclei were excluded during the gating process. 20,000 nuclei were collected in 40 μL resuspension buffer chilled to 4°C to reach the final concentration of 500 nuclei/μL. Sorted nuclei were processed immediately on the 10x Genomics Chromium platform using the Chromium Next GEM Single Cell 3′ Reagent Kit v3.1 according to the manufacturer’s instructions. When fewer than 20,000 nuclei were recovered, all available nuclei were collected in reduced volume and the concentration adjusted. Overall, snRNA-seq data were generated from 27 tumor samples across three longitudinally sampled PDX models (S02714, n = 6; S02906, n = 9; and S02881_S2, n = 9; **Supplementary Table 2**).

All single-cell and single-nuclei sequencing libraries were sequenced on an Illumina NovaSeq 6000 platform. For quality control, libraries were initially sequenced to a depth of 10 million total reads to assess the number of targeted cells, the number of detected genes, and mitochondrial transcript content. Based on the estimated number of high-quality cells, further sequencing of the library was performed to obtain a target depth of 50,000 reads per cell.

#### Processing of scRNA-seq and snRNA-seq data

Single-cell and single-nuclei RNA-seq data were processed using our in-house pipeline *M3K*. Barcodes and unique molecular identifiers (UMIs) were extracted from Read 1, with single-nucleotide barcode mismatches corrected against a reference list and barcodes with multiple mismatches discarded. To account for PDX-derived tumor cells, Read 2 sequences were independently aligned to the human (GRCh37/hg19; hs37d5 decoy) and mouse (GRCm38/mm10) reference genomes using STAR v2.5.2b ^82^. Host-derived cells were identified based on the fraction of human-mapped reads 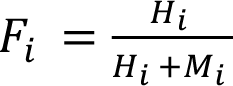 using an inflection-point cutoff to retain cells with at least 75% human-mapped reads. Cell barcodes were then ranked by total read count to generate a barcode rank plot. An automated cut-off was determined at the inflection point and manually inspected to distinguish cell-associated barcodes from low-count background barcodes, which were excluded. Following filtering, genes × cell UMI count matrices were generated for downstream analyses.

Count matrices were loaded separately into *Seurat* v5.1 ^90^. Genes detected in at least three cells and cells with more than 200 detected genes were retained. Cells with >20% mitochondrial reads or abnormal relationships between nFeature_RNA and nCount_RNA, assessed by visual inspection, were excluded. Doublets were identified and removed using *scDblFinder*^91^ after conversion to *SingleCellExperiment*^92^ v1.24.0. Following filtering, single cell RNA-seq of viable tumor cells yielded a total of 369,146 cells from 33 control- and 63 chemotherapy-treated tumor samples across 33 PDX models. Approximately 3,800 cells were retained per sample (median: 4,217; IQR: 2,220–5,222), with a median of 3,620 detected genes (IQR: 3,088–4,146) and 11,170 UMIs (IQR: 8,817–14,414) per cell. For snRNA-seq, performed on longitudinally collected flash-frozen tumor samples, our analysis yielded a total of 82,503 nuclei from 24 tumor samples across three PDX models, representing 19 longitudinal timepoints. A mean of 3,438 nuclei was retained per sample (median: 3,246; IQR: 1,268–4,945), with a median of 952 detected genes (IQR: 638– 1,296) and 1,313 UMIs (IQR: 842–1,908) per nucleus. Consistent with previous reports, snRNA-seq yielded fewer detectable genes and a lower proportion of protein-coding transcripts compared with scRNA-seq^93^. Furthermore, in some samples collected during chemotherapy, fewer nuclei were retained likely reflecting reduced numbers of intact nuclei under chemotherapeutic pressure (**Supplementary Table 9**). For downstream analyses, replicate samples representing the same treatment stage were pooled (**Supplementary Table 2**).

Counts were log-normalized, and the 2,000 most variable genes were identified using the variance-stabilizing transformation (vst) method. Data were centered and scaled while regressing out nCount_RNA, nFeature_RNA, and percent.mt. Cell cycle scores for S and G2/M phases were calculated using *CellCycleScoring* ^94^, followed by re-scaling with additional regression of S.Score and G2M.Score. Principal component analysis (PCA) was performed on variable genes, retaining the minimum number of principal components (PCs) explaining ≥80% of cumulative variance. These PCs were used to construct shared nearest-neighbor (SNN) graphs, followed by Leiden clustering^95^ at a resolution of 0.1 consistently selected for biological interpretability. The same PCs were used for dimensionality reduction by t-distributed stochastic neighbor embedding (t-SNE) or Uniform Manifold Approximation and Projection (UMAP). For each model, Seurat objects from control and chemotherapy-treated samples were merged while preserving cell IDs. Normalization and scaling were repeated, regressing out nCount_RNA, nFeature_RNA, percent.mt, S.Score, and G2M.Score. Scaled data were then quantile-normalized using *normalize.quantiles* from the *preprocessCore* package, and PCA, dimensionality reduction, and clustering were repeated as described above.

To verify sample identity and confirm the presence of patient-derived tumor cells, *peiflyne vartrace* was applied to merged Seurat objects to detect in scRNA-seq reads model-specific somatic SNVs identified by matched WES data. This confirmed identity and presence of tumor cells in all scRNA-seq and snRNA-seq samples. In the scRNA-data from model S03534, contamination from another tumor sample was identified. While uncontaminated baseline WES/WGS data were available for this model, the corresponding tumor sample from the chemotherapy experiment was contaminated and therefore excluded from genomic analyses (**Supplementary Table 2**).

As this contamination referred to only a small fraction of cells in 3′ RNA-seq data, clusters predominantly comprising these contaminant cells were entirely excluded from downstream analyses. These cells and corresponding reads were also removed from the deposited raw sequencing data.

#### Single-cell copy number variation analysis

Copy number variations (CNVs) were inferred from scRNA-seq data using a combined count matrix generated from QC-filtered and merged Seurat objects for all samples from the same patient. CNV calling was performed with our in-house pipeline *sirca*. First, patient-matched WES data were used to generate a unified copy-number segmentation with *Sclust cn*, including shared genomic breakpoints identified by *Sclust ms*^78^. The *sirca cnmap* module was applied to create a copy number (CN) map containing total read counts per gene (pseudobulk) with corresponding copy-number values. To establish an SCLC-specific model, *sirca cnmod* identified genes with expression levels that significantly correlated with CN across multiple SCLC samples (q < 0.001). This model was used by *sirca cncall* to infer CNVs in individual cells using default parameters (minimum of 5 genes per segment, 5 counts per gene, and q-value cutoff of 1×10^-4^). To identify cell populations with similar CNV profiles and reduce single-cell noise, CNV calls (*seg_counts.txt*) were imported into R and visualized using UMAP (*uwot* package). Distinct CNV populations were defined by *k*-means clustering, with the number of clusters determined by their separation in UMAP space. CNV-based clusters were re-analyzed using *sirca cncall* to generate refined subpopulation-level CNV profiles, which were inspected using IGV v2.8.0^80^.

#### Quantification of intra- and inter-sample transcriptional heterogeneity in scRNA-seq data

Transcriptional intratumoral heterogeneity (ITH) and inter-sample heterogeneity were assessed from single-cell transcriptome data using three complementary metrics capturing within-sample transcriptional diversity, global changes in cluster composition and local emergence of treatment-related cell states (**Supplementary Table 10**). 1) To quantify within-sample transcriptional diversity (α-diversity, **Supplementary Fig. 3e**) pairwise Euclidean distances between cells were determined based on the top 2,000 variable genes (*FindVariableFeatures*, *vst*), following normalization and scaling with regression of *nCount_RNA*, *nFeature_RNA*, *percent.mt*, *S.Score*, and *G2M.Score*. The interquartile range (IQR) of these distances served as a measure of transcriptional heterogeneity (adapted from Wu et al.^96^). Mean IQRs per model and condition were compared using paired Wilcoxon signed-rank tests. 2) To determine global inter-sample transcriptional heterogeneity for each model (β-diversity, **Fig. 2c and 2f–g**), we quantified Jensen– Shannon divergence (JSD) between Leiden cluster proportions in control and both chemotherapy-treated samples (chemo 1 and chemo 2, *philentropy* R package^97^ v0.9.0), ranging from 0 (identical distributions) to 1 (maximally different distributions). Jensen–Shannon distance (JSDist), calculated as the square root of JSD, was used for visualization and continuous comparisons and group differences were assessed using Wilcoxon rank-sum tests. Models were classified as exhibiting a transcriptional shift if the maximum JSD across control–chemo comparisons (JSD_max_) exceeded 0.1 or if condition-specific clusters emerged or expanded beyond background variability ^98,99^. 3) We further analyzed local transcriptional turnover (**Fig. 5b**) for each model by identifying chemotherapy-treated cells occupying transcriptional space not shared with control cells using a neighborhood-based approach (adapted from Dann et al.^100^). For each treated cell, the 30 nearest neighbors (*k* = 30) were identified using the first ten principal components (*FNN* v1.1.4.1). Cells with neighbors exclusively present in chemotherapy-treated samples were classified as “novel”, and their percentage was calculated per model. Group differences were assessed using one-sided Kolmogorov–Smirnov tests and Fisher’s exact tests with a descriptive threshold of >15% for “novel” cells.

#### Annotation of transcription factor expression data

Normalized scRNA-seq/snRNA-seq expression values or bulk RNA-seq FPKM/TPM values for *ASCL1*, *NEUROD1*, *POU2F3*, and *YAP1* were used to define thresholds for the expression of SCLC-related transcription factors (TFs). For bulk RNA-seq (PDX models or SCLC cell lines), expression density distributions were computed across all samples, and expression thresholds were set at the first local minimum following the initial peak, separating low from high expression. These cutoffs were used to assign samples to the respective TF subtypes. For PDX models, expression above the threshold for two TFs was classified as combined TF expression (e.g. “*ASCL1*+*NEUROD1*), for cell lines only the dominant TF was annotated. The TF subtype of H211 was annotated based on already published data^101^. For sc-RNA-seq data, we determined for each tumor sample the proportions of cells expressing individual lineage-defining NE TFs, as well as the proportion of cells exhibiting co-expression of multiple NE TFs in individual cells (**Suppl Fig. 3g**). TF expression-defined subtypes were assigned based on normalized expression of *ASCL1*, *NEUROD1*, *POU2F3*, and *YAP1*. For each TF, an expression threshold was determined as the first local minimum following the main peak of the pooled single-cell expression density distribution across all PDX models. Cells were classified as positive for individual TFs or their combinations based on these cutoffs. Cells positive for none of the four TFs were classified as TF-low, while all TF combinations not assigned to a separate subtype were classified as “Other ≥2 TFs”. Subtype proportions were calculated per sample. For statistical comparisons, chemotherapy-treated replicates were averaged per model and compared with matched control samples using a two-sided paired Wilcoxon signed-rank test. Transcription factor assignments based on bulk-RNAseq and sc-RNAseq data are summarized in **Supplementary Table 20**.

#### Tracking of platinum-damaged clones in scRNA-seq data

Single-cell-inferred CN clusters were linked to WES-defined mutation clones based on their relative proportions (**Fig. 4a**, **Supplementary Fig. 7b**). CN clusters concordant with platinum-damaged clones (defined by platinum-associated mutational signatures) were traced in scRNA-seq to characterize their transcriptional programs. In patient-matched chemo-naïve and post-therapy CDX models with acquired platinum-associated signatures (sample S02730), gene-expression programs of untreated control samples were compared to identify processes associated with platinum exposure.

#### Differentially expressed gene (DEG) analysis and gene set enrichment analysis (GSEA) of scRNA-seq data

To identify chemotherapy-associated transcriptional changes, we performed differential expression analysis of platinum-damaged cell populations identified by single-cell tracking (**Fig. 4a, Suppl. Fig. 7c**) or treatment condition-specific Leiden clusters (**Suppl. Fig. 8a–b**). For the latter, condition-specific clusters were identified by comparing their relative proportions between chemotherapy-treated and control samples within each model. Clusters were defined as condition-specific based on the following criteria: (1) <1% in the comparison condition and ≥5% in the tested condition; (2) 1–<5% in the comparison condition, ≥10% in the tested condition, and ≥2-fold enrichment; (3) 5–15% in the comparison condition, ≥2-fold enrichment, and ≥15 percentage-point increase; or (4) >15% in the comparison condition and ≥3-fold enrichment. If specific clusters were detected in only one condition, all cells from the comparison condition were used as reference (**Supplementary Table 10**).

DEGs were identified using Seurat’s *FindMarkers* function with *MAST*^102^ and default parameters. For the analysis of platinum-damaged cell populations, MSigDB Hallmark^53^ enrichment was performed using *fgsea* (v1.28.0) with log₂ fold change-ranked DEGs.

For the broader condition-specific analysis, corresponding cells were pooled across condition-specific clusters within each PDX model and subjected to differential expression analysis. Genes were retained at an FDR-adjusted *P* < 0.01, absolute log₂ fold change > 0.58 (∼1.5-fold), and expression in ≥15% of cells in either group. DEGs were aggregated by response group (“sensitive” or “resistant”) and direction of regulation. Genes were considered recurrently upregulated in chemotherapy-sensitive models if upregulated in 4 models with ≤1 showing downregulation or in ≥5 models with ≤3 showing downregulation. For chemotherapy-resistant models, genes were retained if upregulated in ≥3 models with ≤2 showing downregulation. Gene lists were restricted to protein-coding genes using *biomaRt*^103^, excluding ribosomal and mitochondrial genes.

For exploratory functional characterization, recurrently upregulated genes were classified using a heuristic approach based on MSigDB Hallmark^53^, Reactome^104^, and GO slim^105,106^ annotations. Annotation terms were assigned by keyword matching to ten predefined biological categories, comprising immune/inflammation, metabolism/hypoxia, EMT/migration/ECM/tissue integrity, cell death/p53/cellular stress, translation/protein homeostasis, chromatin/transcriptional regulation, development/signaling/hormone, cell cycle/proliferation/mitosis, NOTCH/neuroendocrine/neuronal/cell fate, and DNA repair/genome stability (**Supplementary Table 16**). Category scores were calculated from weighted contributions of Hallmark (0.50), Reactome (0.25), and GO slim (0.25), normalized by available annotation sources. Individual terms were additionally weighted according to biological specificity, prioritizing direct functional evidence (2.5) and selected Hallmark pathways (2.0) over standard (1.0) or broad/context-dependent annotations (0.35). To avoid redundancy, only the strongest matching term per annotation source and category was retained. Each gene was assigned a primary category among those scoring within 0.1 of the highest score, prioritizing direct functional evidence, the number of supporting annotation sources, and category score, with category frequency used as the final tie-breaker. Genes without matching annotations were classified as “Other”

#### Gene signature scoring in scRNA-seq data

Gene signatures were obtained from published studies, including mesenchymal and epithelial signatures^55^, basal and neuroendocrine (NE) signatures^76^, and a YAP/TAZ target signature^77^, as well as MSigDB Hallmark^53^ gene sets. Additional custom gene signatures were derived from the recurrently upregulated genes assigned to functional categories as described above. Signature activity was quantified using *Seurat’s AddModuleScore* function, which calculates average signature gene expression relative to expression-matched control genes. For the Tan et al.^55^ signatures, comprising predefined sets of epithelial and mesenchymal marker genes, epithelial and mesenchymal module scores were calculated separately for each cell. A continuous EMT score was then calculated as the difference between the mesenchymal and epithelial module scores (mesenchymal − epithelial), with higher scores indicating a more mesenchymal transcriptional state. Epithelial and mesenchymal module scores and the derived EMT score were visualized on UMAP embeddings.

#### CellOT analysis

To complement differential gene expression analyses, we applied CellOT, a neural optimal transport framework for predicting cellular responses to perturbations from unpaired single-cell data^56^ . Rather than focusing on individual genes or predefined cell populations, we applied CellOT to pathway activity scores derived from scRNA-seq data to characterize chemotherapy-associated shifts in transcriptional programs. The analysis was restricted to sensitive and resistant models, excluding intermediate responders.

Cells with at least 1,000 counts were retained, and expression data were normalized to 10,000 counts per cell and log-transformed using Scanpy^107^ (v1.10.4). Count matrices were aligned to a common set of genes, with missing genes assigned zero expression.

Two feature sets were analyzed independently: (i) the 50 MSigDB Hallmark gene sets^53^ combined with epithelial and mesenchymal signatures from Tan et al.^55^ and (ii) the epithelial and mesenchymal signatures from Tan et al.^55^ alone.

Pathway activity scores were calculated jointly across sensitive and resistant cells using gene set variation analysis (GSVA)^108^ implemented in GSEApy^109^ (v1.1.7) and subsequently Z-score standardized per pathway.

For each feature set, separate CellOT models were trained for chemotherapy-sensitive and - resistant tumors to map control cells to predicted chemotherapy-treated states. Cells were randomly divided into training (80%) and validation (20%) sets. CellOT models were trained with four hidden layers of 64 units each, a batch size of 64, and the Adam optimizer^110^ (learning rate = 10⁻⁴, β₁ = 0.5, β₂ = 0.9, weight decay = 0) for 50,000 iterations. Model performance was evaluated every 250 iterations using maximum mean discrepancy (MMD) between predicted and observed treated-cell distributions, and the checkpoint with the lowest validation MMD was retained. The retained map was applied to all control cells to predict their chemotherapy-treated states, with cell-level pathway shifts (Δ) calculated as the difference between predicted treated and original control scores (Δ = predicted treated − control).

To visualize treatment-associated shifts in transcriptional programs, principal component analysis (PCA) was fitted to Z-score standardized pathway activity scores from all observed control and chemotherapy-treated cells. Control cells and their CellOT-predicted chemotherapy-treated counterparts were subsequently projected into this PCA space. Chemotherapy-associated shifts were calculated as the differences between the coordinate-wise median positions of control and predicted cells on PC1 and PC2 (**Fig. 5c**). Shift magnitude was quantified as the Euclidean distance between these positions, while shift coordination was assessed using cosine similarity between each model’s shift vector and the median shift direction of its response group (**Suppl. Fig. 8f**). Pathway-specific cell-level shifts Δ*_im_*_g_ were analyzed using linear mixed-effects models (LMMs) to account for dependence among cells from the same tumor model: Δ*_im_*_g_ = *β_r_*_g_ + b_mg_ + ε_img_, where Δ_img_ represents the pathway-specific (g) shift for a cell (i) from model (m), *β_r_*_g_ the mean shift for response group (r), and b_mg_ the model-specific random intercept. Models were fitted by restricted maximum likelihood with Powell optimization using Statsmodels^111^ (v0.14.4). Response group-level effects (*β_r_*_g_; **Fig. 5e, Suppl. Fig. 8h**) were assessed using two-sided Wald tests, with Benjamini–Hochberg correction across pathways within each response group. For the focused EMT analysis (**Fig. 5f**), model-specific effects were calculated as 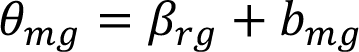, with standard errors estimated as: 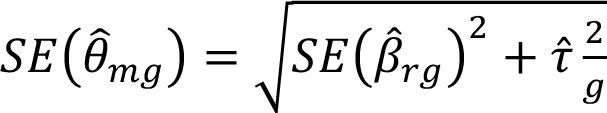 where 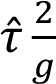 denotes the estimated between-model variance. Two-sided normal-approximation P values were adjusted across the two signatures within each model using Benjamini–Hochberg correction with q ≤ 0.05 considered significant.

#### Cell culture of established cell lines

Established, immortalized human SCLC cell lines were primarily obtained from ATCC, and some were provided by academic collaborators. Their identity was verified by STR profiling, and all cell lines were routinely tested for the absence of mycoplasma. Cell lines were cultured in RPMI (Gibco), DMEM (Gibco) or HITES medium based on DMEM/F12 (Thermo Fisher Scientific) containing 10 nM hydrocortisone, 0.005 mg/ml insulin, 0.01 mg/ml transferrin, 10 nM β-estradiol and 30 nM sodium selenite supplemented with 1% L-glutamine, 1% penicillin/streptomycin cocktail and 10% FBS. Cells were maintained in a humidified atmosphere at 37 °C with 5% CO₂.

#### CTG viability assay

Cell viability after treatment with cisplatin or etoposide (Central Cytostatic Preparation Unit, Pharmacy of the University Hospital Cologne) was assessed after 96 h using the CellTiter-Glo reagent (Promega #G7570) and normalized to the DMSO-treated controls. Cells were seeded at cell line specific densities of 500–10,000 cells into flat-white 96-well plates (Corning) and treated with nine 3-fold serial dilutions (0.0045–30 μM) or DMSO after 24 h. Half-maximal inhibitory concentrations (IC_50_) were determined by fitting sigmoidal four-parameter logistic (4PL) dose– response curves using nonlinear regression and interpolation at 50% viability in R (*minpack.lm*, *nlsLM*). SCLC cell lines were classified as sensitive or resistant if both cisplatin and etoposide IC_50_ values were below or above their respective medians, respectively. Data are presented as mean ± SEM, and statistical significance was assessed using Welch’s t-tests on log_10_-transformed IC_50_ values.

#### Generation of chemotherapy-resistant cell lines *in vitro*

Chemotherapy-sensitive human SCLC cell lines (H1048, GLC1, H526, H1694) were selected to induce chemotherapy resistance by continuous exposure to stepwise escalating concentrations of either etoposide or cisplatin (prepared by Central Cytostatic Preparation Unit, Pharmacy of the University Hospital Cologne) concentrations over a period at least 6-8 months. Media supplemented with chemotherapeutic agents was replenished at least 2 times per week. During this period, cellular responses were monitored by CTG viability dose-response assays; and drug concentrations were progressively increased until IC_50_ values were 5–10-fold higher than in the respective parental cell line. Parental cells were maintained and passaged in parallel to control for potential effects of long-term cultivation. For GLC1, the resistance experiment was performed twice independently, generating biological replicates annotated as “GLC1 #1” and “GLC1 #2”. Detailed information on the cell lines, experimental conditions, and collected samples are provided in **Supplementary Tables 13 and 14**.

#### Visualization software and statistics

Plots for data visualization were generated in R using *ggplot2* (v3.5.2) or GraphPad Prism (v10.1.1). The oncoprint visualization was created using the *ComplexHeatmap* (v2.18.0) package in R. Figures were further adjusted using Adobe Illustrator (v29.7.1). Statistical analyses were conducted with GraphPad Prism or R (v4.3.2), with test selection based on data type and experimental design. Details of the applied statistical tests are provided in the corresponding figure legends.

## Acknowledgements

We sincerely thank all patients who contributed tumor specimens to this study and their families for their support. We are also grateful to the physicians, nurses, and other clinical staff involved in patient care and sample collection, whose efforts made this study possible. We thank current and former members of the Peifer, George, and Thomas laboratories for their experimental and computational support, as well as valuable discussions throughout this study. This work was supported by the German Research Foundation (DFG) through funding provided by a collaborative research center grant on small cell lung cancer (CRC1399, project-ID 413326622) to JG, RKT, MC, MP, KH, RB, AQ, LATF and LW. JG, RKT, and MP received funding from the German Federal Ministry of Education and Research (BMBF, e:Med consortium InCa, grants 01ZX1901A and 01ZX2201A). RKT was supported by the European Regional Development Fund (EFRE) initiative of the German state of North Rhine-Westphalia (EFRE-0800397). Additional funding was received from “Netzwerke 2021”, an initiative of the Ministry of Culture and Science of the German state of North Rhine-Westphalia, for the CANTAR project to JG, RKT, and MP. Further support was provided by the DFG (project ID: 497777992 to JG, MP, and AQ; project ID: GRK3110 to JG, MP and RB; project ID: 418074181 to MP; and through CRC 1530, project ID: 455784452, to MP), the German Cancer Aid (TACTIC, project ID 70115201, to RKT, and Mildred Scheel Professorship to MP), the Bruno-Helene-Joester Foundation (JG and MP), and the Jean Uhrmacher Foundation (JG). GG was supported by the Barbara Massie Memorial Fund, the MDACC Moonshot FIT Program, the Bruce Krier Endowment Fund and the Lyda Hill Foundation, the NIH/NCI (CA285454-01A1, CA288448-01), the Department of Defense (KC200096), the NIH (CA258226-01) and the CPRIT-IIRA RP260133. LZ is supported by National Institutes of Health National Cancer Institute K99/R00 (1K99CA307136-01). WJ and KVL were funded under the Excellence Strategy of the Federal Government and the Länder. We thank the Cologne Center for Genomics (CCG), part of the West German Genome Center (WGGC), for their assistance with library preparation, sequencing, data processing and STR analysis. We also acknowledge the German Cancer Research Center (DKFZ) for performing single-cell RNA sequencing, supported by funding from the German Cancer Consortium (DKTK) to RKT. We thank the FACS & Imaging Core Facility at the Max Planck Institute for Biology of Ageing for their support with cell sorting by flow cytometry. We are also grateful to the staff of the Animal Husbandry Network (THNW) of the Faculty of Medicine at the University of Cologne for their dedicated animal care and support throughout this study. We furthermore thank the IT Center University of Cologne (ITCC) for providing computational resources on the DFG-funded high-performance computing clusters CHEOPS and RAMSES, as well as technical support (DFG funding number: INST 216/512-1 FUGG).

## Author contributions

LCK, RKT and JG conceived and designed the study. LCK performed experiments and data analyses, with experimental and technical support from CM, NA, CL and OS. WJ and KVL developed and implemented the CellOT-based analytical approach. LM, MN, AR and MP developed and maintained computational pipelines and workflows. LM, MN, AR, MS, MC and DP contributed to computational and statistical analyses. WJ, LZ, LATF, GG, KVL and MP provided scientific input, performed data analyses and contributed to data interpretation. FJ, KH, JPW, BH, FD, JPP and LH contributed patient samples, supported collection of clinical data, and performed clinical data analysis. CPA and LW supported sample processing and *in vivo* experiments in mice. MG and ALS performed nuclei sorting. AQ and RB performed histopathological evaluations. MB and CC performed STR profiling. AM supported genomic data deposition. GB coordinated project administration and research resources and provided logistical support. RKT and JG supervised the study. LCK and JG wrote the manuscript. WJ, LZ, LATF, GG, MC, MP and RKT critically revised the manuscript for important intellectual content. All authors reviewed and approved the final manuscript.

